# Differential effects of soil conservation practices on arthropods and crop yields

**DOI:** 10.1101/2021.12.06.471474

**Authors:** Elinor M. Lichtenberg, Ivan Milosavljević, Alistair J. Campbell, David W. Crowder

## Abstract

Many agricultural management tactics, such as reduced tillage, aim to promote biodiversity and ecosystem services. Responses to such tactics can be context dependent, however, and differentially impact (i) functional groups of service-providing organisms and (ii) crop yields. In canola (*Brassica napus* L., *B. rapa* L.) crop fields we assessed how soil tillage and landscape context (amount of semi-natural habitat within 1 km of each field) affected arthropod biodiversity and crop yield. We assessed effects of full (multiple tillage passes that leave soil surface bare), intermediate (tilled once and some stubble remains), or no (seed planted directly into last year’s stubble) tillage on functional groups with unique diets and reproductive strategies: (i) herbivores, (ii) kleptoparasites, (iii) parasitoids, (iv) pollinators, and (v) predators. Effects of tillage and landscape context on arthropod abundance and diversity varied across functional groups. Pollinators responded strongest to tillage, benefitting from intermediate tillage. Predators and herbivores responded strongly to landscape context, as both were more abundant in landscapes with more semi-natural habitat. Our results suggest natural history differences among functional groups mediate effects of landscape context on biodiversity. However, variation in arthropod communities had little effect on canola crop yield. The effects of soil management practices on aboveground arthropods are complex, and practices thought to increase some aspects of agricultural sustainability may not be beneficial in other contexts. Identifying practices such as intermediate tillage that may increase soil quality and arthropod diversity is a key to designing agricultural ecosystems that will effectively benefit both biodiversity and human well-being.

## Introduction

Agricultural management tactics such as intercropping and reduced tillage are implemented in an effort to support biodiversity and ecological services in agroecosystems without sacrificing crop yield. Reduced tillage, for example, supports biodiversity by creating soil habitat availability and heterogeneity (de Graaff, Hornslein, Throop, Kardol, & van Diepen, 2019; Tamburini et al., 2020). However, responses of organisms to practices such as tillage often vary among service-providing functional groups (Lefcheck et al., 2015; Mitchell et al., 2015). This has led to calls to better assess impacts of habitat change on multiple ecosystem services such as soil quality, pollination, and crop yield (Bommarco, Kleijn, & Potts, 2013; Tamburini et al., 2020). Given the complexity of agricultural food webs, studies should also assess how organisms in unique functional groups respond to management, and whether there are trade-offs or synergies in managing farms for different ecosystem services.

Effects of in-field soil management practices on ecosystem service providers likely depend on how animal functional groups interact with soil. For example, tillage can harm ground-nesting bees (Ullmann, Meisner, & Williams, 2016) and predators that shelter among weeds (Cranshaw, 2004), but often has little to no impact on herbivorous pests that feed on crops (Tooker, O’Neal, & Rodriguez-Saona, 2020). Reduced tillage is implemented to limit soil erosion and conserve soil moisture. However, it can also affect soil chemical and physical profiles, and availability of flowering plants that feed beneficial insects (A. C. Kennedy & Schillinger, 2006). Impacts of soil management practices on distinct functional groups may also depend on the landscape context. Semi-natural habitat near farms can facilitate dispersal of organisms to crops (Kremen et al., 2007; Tscharntke et al., 2012) and responses of organisms to particular practices may depend on the landscape context. Indeed, some studies show benefits of sustainable agricultural practices accrue most strongly in relatively simple landscapes, while others show the greatest benefits in complex landscapes (C. M. Kennedy et al., 2013; Lichtenberg et al., 2017; Scheper et al., 2013; Tscharntke et al., 2012).

While supporting biodiversity is often a goal of agricultural production systems, crops must also generate high yields. Because yield captures the total contribution of biotic communities and farming practices, it can be difficult to ascribe yield to individual factors (Tamburini, Bommarco, Kleijn, van der Putten, & Marini, 2019). Studies are thus needed that use a single analytical framework to assess how habitat availability and diversification affect biodiversity and yield, both directly and through indirect interactions among organisms and management practices (Birkhofer et al., 2015; Byrnes et al., 2014; Weekers et al., 2022). Such studies may be particularly useful when conducted in fields that involve commercial production and representative growing practices for a given region.

Here we assessed how soil tillage affected arthropod functional groups in canola (*Brassica napus* L., *B. rapa* L.) crops of the Pacific Northwest USA, and how landscape context and tillage interacted with arthropods to affect yields. Canola provides floral food for pollinators and natural enemies, but attract pests like aphids (Aphididae) and flea beetles (*Phyllotreta cruciferae* Goeze [Chrysomelidae]). We first asked if effects of tillage on arthropod abundance and diversity varied by functional group. We hypothesised functional groups with soil dependence, like pollinators and kleptoparasites, would be most strongly negatively impacted by tillage (Rowen, Regan, Barbercheck, & Tooker, 2020). Second, we asked if responses of functional groups to tillage depended on landscape context, given variation in mobility and habitat needs of unique organisms (Lichtenberg et al., 2017; Marja, Tscharntke, & Batáry, 2022). Third, we asked whether variation in arthropod communities interacted with tillage and landscape context to affect yield (Delaplane & Mayer, 2000; Morandin & Winston, 2005; Reddy, 2017). Reduced tillage often decreases crop yield (e.g., Lundin, 2019; Tamburini et al., 2020), but enhanced pest control or pollination could counteract this. This allowed us to assess how agricultural practices directly and indirectly affected multiple ecosystem services and biodiversity across landscapes.

## Materials and Methods

### Study sites

We sampled arthropods in spring canola fields in eastern Washington and northern Idaho during 2013 and 2014 (Fig. S1; Table S1). This heavily agricultural region has patches of semi-natural habitat amidst considerable acreage of grains, legumes, and canola (Painter, Hinman, & Roe, 2006; USDA NASS, 2020). The region’s loess soils are sandy loams. As a mass-flowering crop that blooms for up to a month, canola attracts flower-feeding arthropods such as pollinators and natural enemies like predators and parasitoids (Delaplane & Mayer, 2000; Morandin & Winston, 2005). Canola fields are thus an effective model system for studying multiple ecosystem service providers across unique functional groups.

We selected 15 spring-planted canola fields each year ranging from 0.7 to 142 ha (mean ± SD = 44.4 ± 41.0). The short canola bloom period and relatively small number of spring canola fields in the region limited further sampling. Most fields were maintained by local farmers with four maintained by local universities or seed companies. Farmers used several different canola varieties, sometimes within one field. Most of these varieties were glyphosate resistant, with one farmer using Beyond^®^ (herbicide) resistant seed. All seeds were treated with a neonicotinoid (mainly thiamethoxam), and some sites applied a pyrethroid insecticide once after bloom to control flea beetles and cabbage seed pod weevils (*Ceutorhynchus obstrictus* [Marsham]). Most farmers also applied a herbicide treatment (glyphosate) before bloom. Informal conversations with farmers indicated that canola variety used, and glyphosate and pyrethroid application, were independent of tillage regime. These canola fields were typically immediately adjacent to grain (barley, wheat) or legume (garbanzo beans, lentils, peas) crops, with little non-crop vegetation along field edges. Our canola fields were spaced at least 2 km apart, which was sufficiently far relative to insect flight distances to consider them spatially independent.

### Arthropod sampling

We used two collection techniques to sample diverse arthropods that associate with canola: (i) traps typically used to sample bees and (ii) sweep nets. All sampling occurred along one field edge on days with daytime temperatures above 13 °C and wind speed below 4.5 m/s, while fields were in full bloom. Sampling locations were haphazardly selected along accessible field edges, where we could work without damaging crops and where traps on the ground would not be covered in vegetation. At 10:00, we conducted 100 continuous sweeps in the canola canopy. We walked at a steady pace and thus covered approximately the same distance at each field. We then emptied net contents – including all arthropods – into a plastic bag and stored them on ice. In the lab, we freeze-killed arthropods, then sorted them to morphospecies. Bees were pinned and specimens in other taxa were stored in ethanol.

Around 12:00, we set out a line of bee traps that stayed in place for 24 h. This line included two blue vane traps (SpringStar) and six 96.1mL pan traps (7 cm diameter) painted in colours that attract bees (two each of white, fluorescent yellow, fluorescent blue; Kearns & Inouye, 1993; Leong & Thorp, 1999) and partially filled with soapy water. We separated traps by 5 m and located them ∼0.5 m from the field edge. Traps and sweep netting began at the same spot. The following day, we collected trapped arthropods (bees as well as other arthropods) using a strainer, washed off excess soap, and stored the specimens in 95% ethanol. In the lab we washed and dried bees, and sorted all specimens to genus and morphospecies based on morphological characteristics (Arnett, 2000; Arnett & Thomas, 2000; Arnett, Thomas, Skelley, & Frank, 2002; Boyle & Philogène, 1983; Derraik et al., 2002; Derraik, Early, Closs, & Dickinson, 2010; Michener, 2000; Stehr, 1987a, 1987b). Both adults and immature forms of all arthropod groups were considered.

Specimens were also assigned to five functional groups using literature and information from local species (Arnett, 2000; Michener, 2000; Stehr, 1987a, 1987b): (i) pollinators, (ii) herbivores, (iii) predators, (iv) parasitoids, and (v) kleptoparasites (Table S2). This classification considered both diet and reproductive strategy. Herbivores included major regional canola pests (thrips [Thysanoptera], aphids, flea beetles, cabbage seedpod weevils, and *Lygus* bugs Hahn; Reddy, 2017). Kleptoparasites are regulators of bee communities, and may reflect overall levels of pollinator biodiversity (Sheffield, Pindar, Packer, & Kevan, 2013). We also considered predators and parasitoids pooled together as “natural enemies”.

### Field and landscape variables

For each site we assessed agronomic and weather factors that can affect insects: (i) tillage, (ii) field size, (iii) growing degree days, and (iv) cumulative precipitation (e.g., Aldercotte, Simpson, & Winfree, 2022; Forcella et al., 2021; Fragoso, Jiang, Clayton, & Brunet, 2021; Skellern, Welham, Watts, & Cook, 2017; O. M. Smith et al., 2020). We asked farmers directly to categorize each site’s tillage regime. A field had either (i) full (4 to 7 passes producing bare soil with no stubble before seeding; n = 6), (ii) intermediate (field tilled once with the same equipment as full tillage fields, leaving some stubble remaining before seeding; n = 13), or (iii) no (soil not tilled and seed planted directly into the previous year’s stubble; n = 11) tillage. These represent the three main tillage regimes used in the region. All fields were tilled in spring (typically April into early May). Because canola is planted very shallow, spring tillage in the region is recommended to be limited to 2 to 5 cm (Brown, Davis, Lauver, & Wysocki, 2009). We determined field size in ArcGIS by creating a polygon tracing the boundary and subtracting the areas of all uncultivated patches (typically remnant prairie within and adjacent to fields; originally mapped by hand). We gathered daily temperature and precipitation values from PRISM (PRISM Climate Group, 2020) and calculated growing degree days and cumulative precipitation through the day before we sampled arthropods. These environmental measures can cause inter-annual variation in arthropod populations (Forcella et al., 2021; Skellern et al., 2017) and helped control for mild temperature and precipitation gradients across our study region. PRISM estimates climatic conditions at a given location on a given day (at 4-km resolution) using climatologically-aided interpolation based on 30-year patterns (PRISM Climate Group, 2020). Growing degree days were calculated as the cumulative amount of heat above 5 °C since January 1 (Dickson, 2014). We calculated precipitation cumulatively from January 1.

We also calculated the amount of semi-natural habitat within 1 km of each field, a radius that is ecologically relevant for pollinators, herbivores, and natural enemies (Greenleaf, Williams, Winfree, & Kremen, 2007; Rusch et al., 2016). We determined land cover using CropScape data (USDA SARS, 2014a, 2014b, 2015a, 2015b) to calculate area of each land cover type around each site. Semi-natural habitat included forests, grasslands, shrublands, and wetlands, while crop habitats were classified as agricultural.

### Data analysis: Effects of tillage and landscape on arthropod communities

We used model selection to test how functional groups responded to tillage and landscape context. We calculated three community metrics – (i) abundance, (ii) richness, and (iii) evenness (E_var_, B. Smith & Wilson, 1996) – for each functional group at each site. Evenness captures how individuals are distributed across taxa and indicates the degree that rare taxa affect ecosystem functioning (Crowder, Northfield, Strand, & Snyder, 2010; Winfree, W. Fox, Williams, Reilly, & Cariveau, 2015). Although richness and evenness are more abstract when not all taxa can be identified to species, we consistently identified to morphospecies across all fields and thus our richness and evenness values provide important measures of the sampled communities. We ran linear regressions for each functional group with tillage, proportion semi-natural habitat, their interaction, year, degree days, cumulative precipitation, and field size as fixed effects. These variables were not collinear (Table S3). As abundance data were overdispersed, we used negative binomial regressions (MASS package, Venables & Ripley, 2002). We analysed richness and evenness with linear models (identity link), but herbivore evenness was square-root-transformed due to heteroscedasticity. All analyses met model assumptions. We used information-theoretic model selection to assess model fit for functional group metrics (MuMIn package, Barton, 2014), and selected models with AICc values within 2 of the lowest value (Burnham & Anderson, 1998). While tillage and field size were not collinear (Table S3), fields experiencing full tillage were mostly smaller than fields with no tillage (Fig. S2; Kruskal-Wallis rank sum test: χ^2^_2_ = 10.11, P = 0.006; Dunn’s test of multiple comparisons with a Holm adjustment: no vs. intermediate tillage Z = 2.15, P = 0.03 [not significant with adjustment], no vs. full tillage Z = −3.18, P = 0.002, intermediate vs. full tillage Z = −1.35, P = 0.08; Dinno, 2017; Holm, 1979). We thus conducted model selection first with only the requirement to include year in all models, and then with an additional constraint to include field size if a model retained tillage. Results were the same with and without this constraint (Tables S4-6). We thus present only unconstrained models below. Low richness, and thus lack of variation, prevented us from analysing kleptoparasite and parasitoid evenness; all other metrics were assessed. There were 13 total models, one for each functional group and community metric, each with 30 observations (28 for predator evenness).

We used multivariate PERMANOVA to test whether community composition responded to tillage and landscape context (vegan package, Oksanen et al., 2019). We categorized the following predictors as “high” (above the median) and “low” (below the median): proportion semi-natural habitat, degree days, cumulative precipitation, and field size. Analyses also included tillage, the interaction between tillage and proportion semi-natural habitat, and year as predictors. We used Sørensen dissimilarity indices based on presence/absence matrices with singletons species removed. We analysed each functional group separately. Small numbers of kleptoparasite and parasitoid taxa prevented analysing these groups. To assess each term’s significance via type II sums of squares (as in regressions) we first ran each model with the option for marginal effects. We then tested significance of tillage and the proportion of semi-natural habitat via two additional regressions that used sequential order with the variable of interest assessed after all other fixed effects had been accounted for. To visualize potential community differences as functions of tillage and landscape context, we plotted a separate Principle Coordinate Analysis for each variable and functional group (Oksanen et al., 2019).

### Data analysis: Effects of tillage, landscape and arthropod communities on crop yields

We assessed how arthropod biodiversity, tillage, and landscape context affected canola crop yield;. Farmers provided yield data for their sites directly. We ran separate linear regressions (identity link) with abundance, richness, or evenness that also included tillage, proportion semi-natural habitat, and their interaction, year, degree days, cumulative precipitation, and field size as fixed effects. This enabled us to investigate both direct impacts of tillage and landscape, and direct impacts of arthropods while controlling for tillage and landscape.

Considering these results together with direct impacts of tillage and landscape on arthropod communities enabled us to investigate indirect impacts of habitat on yield via arthropod communities. For each set of models, we selected the best fit models as those with AICc values within 2 of the smallest value. Variables in best models were not collinear (Table S7). As with arthropod community models, we ran models that also included field size and constrained model selection to include field size if a model retained tillage. Results were the same with and without this constraint (Table S8). There were three yield models, one for each community metric, each with 30 observations (28 for the evenness regression).

## Results

We collected 21,446 individuals across 130 taxa, with 15,256 herbivores (20 taxa, 82% of individuals were pest species), 154 kleptoparasites (8 taxa), 1,080 parasitoids (6 taxa), 4,217 pollinators (65 taxa), and 739 predators (32 taxa). Our data included 80 spiders (order Araneae); all other individuals were insects. The most abundant herbivores were thrips, aphids, sciaroid flies, and chrysomelids (7,184, 4,078, 2,041, and 576 individuals, respectively). The most common pollinators were two halictid bee morphospecies (813 *Lasioglossum* and 581 *Agapostemon* individuals). The most common natural enemies were chalcidoid wasps (727 individuals, parasitoid) and melyrid beetles (229 individuals, predator). Landscapes around our sites ranged from 0% to 36% semi-natural habitat. This variable was independent of tillage regime (Kruskal-Wallis test: X^2^_2_ = 2.47, *P* = 0.29).

### Effects of tillage and landscape on arthropod communities

Overall, pollinators and kleptoparasites were affected by tillage while herbivores and predators responded most strongly to landscape context. Pollinator richness was higher in fields with intermediate tillage than fields with no tillage (Fig. 1a; Tables S5, S9). Kleptoparasite abundance was higher in fields with intermediate or no tillage than in heavily-tilled fields in one of four best models (and significant at α < 0.10 in a second model; Fig. 1b; Tables S4, S9). Pollinator abundance and evenness, and kleptoparasite richness, were not affected by tillage or landscape context (Tables S4-6). In contrast, more semi-natural habitat promoted herbivore and predator abundance, and reduced predator evenness, regardless of tillage regime (Fig. 2; Tables S4, S6). Herbivore diversity and predator richness were unaffected by tillage or landscape context (Tables S5, S6). Parasitoids were unaffected by tillage or landscape context (Tables S4-6). Tillage and landscape context never had an interactive effect. They also did not affect community composition of any functional group (Table S10, Fig. S3).

**Figure 1:**
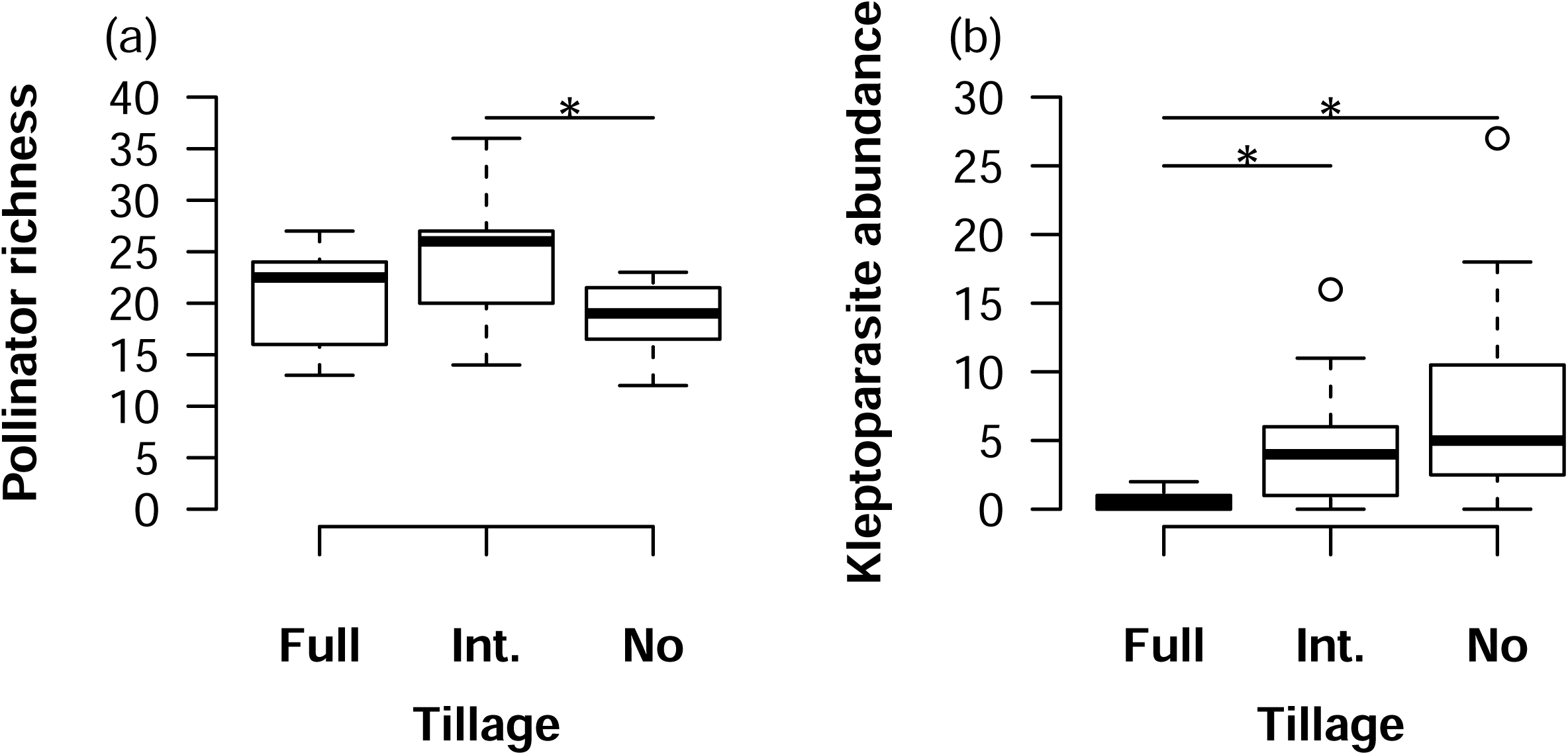
Box and whisker plots of (a) pollinator richness (total # taxa sampled at each site across all sampling methods) and (b) kleptoparasite abundance (total # individuals sampled at each site across all sampling methods) as a function of tillage regime. “Int.” is intermediate tillage. Boxes span the 25^th^ to 75^th^ percentiles, whiskers extend 1.5 times the interquartile range beyond the boxes, and there is a line at the median. Lines with asterisks indicate groups that are statistically different from each other at α = 0.05.

**Figure 2:**
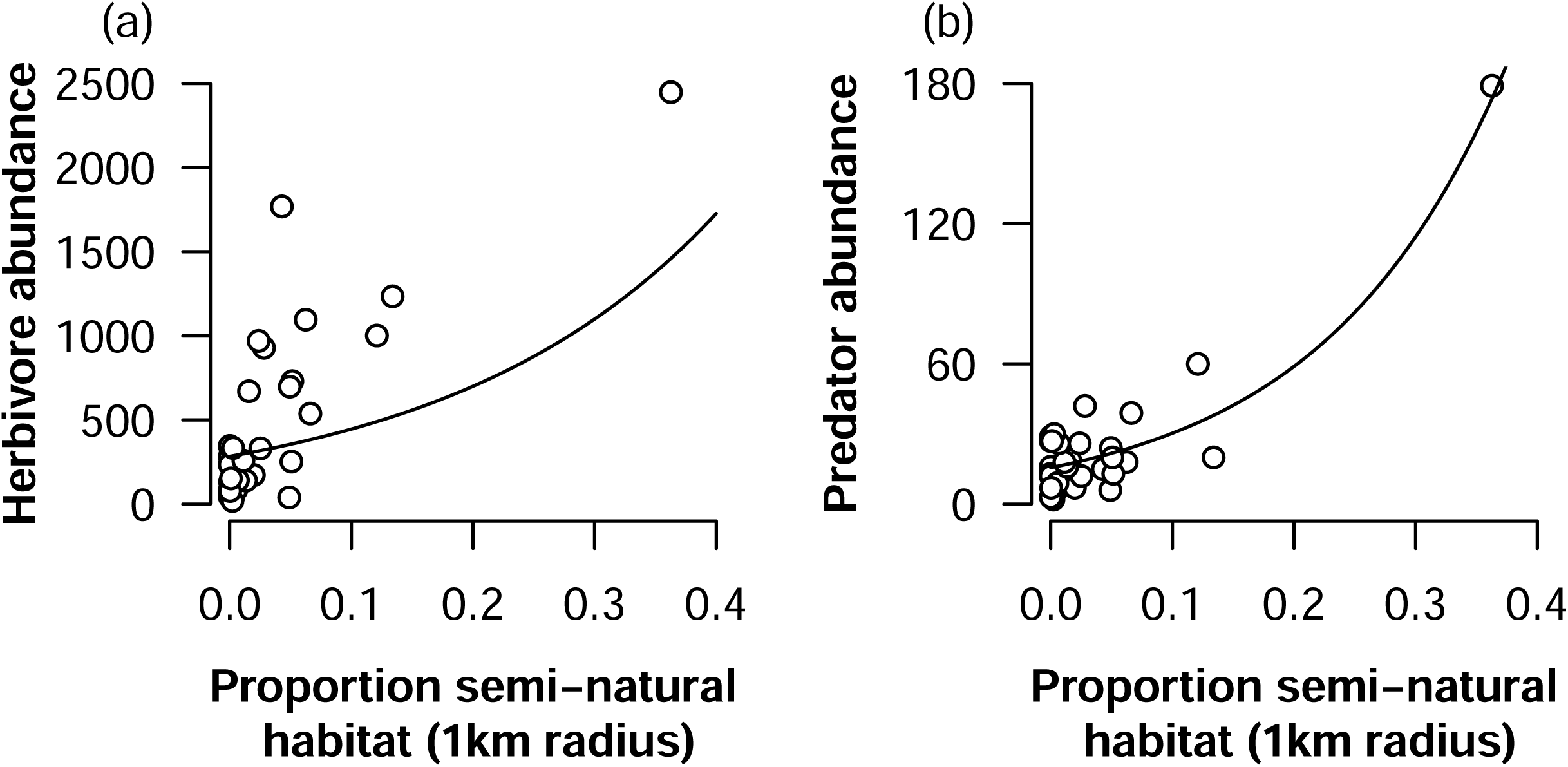
(a) Herbivore and (b) predator abundance (total # individuals sampled at each site across all sampling methods) increase with landscape-scale habitat availability (proportion of semi-natural habitat in a 1 km radius around a site). Curves show best-fit lines from negative binomial regressions.

Environmental variables had mixed effects on arthropod abundance and diversity (Tables S4-6). Precipitation generally had stronger impacts on arthropod communities than degree days. Greater kleptoparasite abundance and richness, pollinator abundance, and predator richness occurred at sites with lower precipitation. Parasitoid abundance and pollinator evenness had the opposite pattern. Kleptoparasite richness, parasitoid abundance, and predator richness were highest at cooler sites. Larger fields were associated with greater kleptoparasite abundance and richness but lower pollinator evenness.

Differential responses of arthropod functional groups may indicate trade-offs when aiming to manage biodiversity, but we found stronger evidence for biodiversity synergies than trade-offs. Functional groups with known trophic relationships often had correlated metrics (Table S11). For example, we found positive correlations in abundance and diversity between pollinators and kleptoparasites, and among parasitoids, predators, and herbivores. Abundance and evenness of pollinators was also positively correlated with abundance and evenness of natural enemies, as well as individual natural enemy groups (predators or parasitoids) (Table S11). However, biodiversity synergies may be agronomic trade-offs. Abundances of beneficial pollinators and detrimental herbivores also positively correlated (Table S11).

### Effects of tillage, landscape and arthropod communities on crop yields

Tillage strongly affected canola yield, and yield was lower in fields with no tillage than with full or intermediate tillage (Fig. 3a, Tables S8 S9). Landscape context did not impact canola yield (Table S8). We found only mild evidence for effects of beneficial arthropods on yield (Table S8). Pollinators and predators were rarely retained in best models, and their effects were never significant when they were retained. Parasitoid richness was retained in two of three best models, and sites with more parasitoid taxa had higher canola yield in one of these models (with *P* = 0.058 in the other). However, more diverse herbivore communities lowered yield (Fig. 3b; Table 8). Arthropod abundance and evenness did not impact yield.

**Figure 3:**
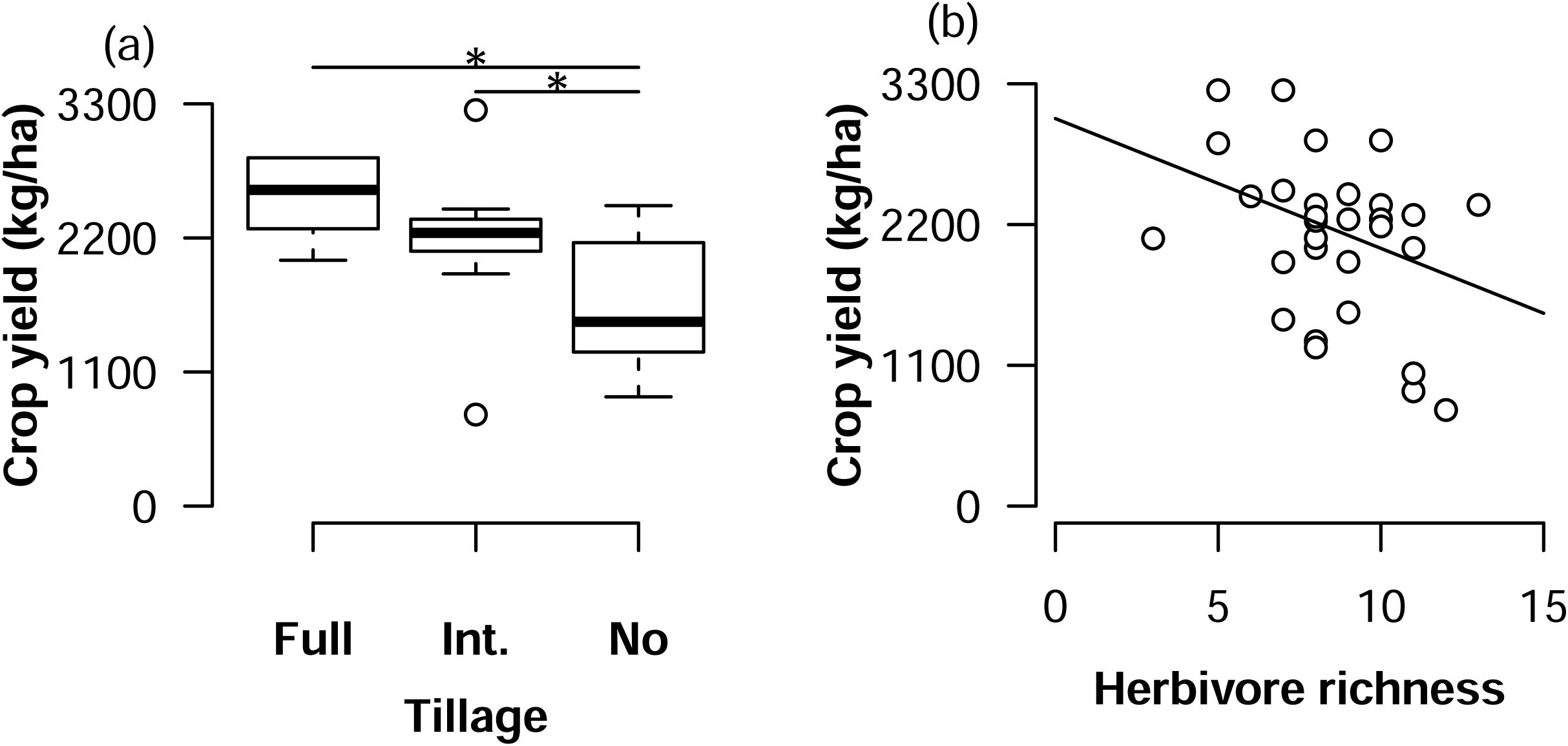
Canola yield (a) was lowest with no tillage and (b) decreased as herbivore richness increased. “Int.” is intermediate tillage. Boxes span the 25^th^ to 75^th^ percentiles, whiskers extend 1.5 times the interquartile range beyond the boxes, and there is a line at the median. Lines with asterisks (a) indicate groups that are statistically different from each other at α = 0.05. The curve (b) shows the best-fit line from linear regression.

## Discussion

Soil management may differentially affect functional groups due to differences in resource needs or dispersal among taxa (Bommarco et al., 2013; Harmon-Threatt, 2020). Because many of these arthropods provide ecosystem services or are pests, biodiversity can affect crop yield. We found that agricultural landscapes can simultaneously support pollinators and predators, but different functional groups respond to habitat variability at different scales. Pollinators were most affected by tillage within fields, while landscape context most strongly affected herbivores and predators. Yet, only herbivores strongly impacted crop yield. Our results showed that agriculture practices alter crop yield directly, but not always indirectly by affecting arthropods (as in Ricketts et al., 2016).

We found that pollinator and kleptoparasite, but not herbivore, predator, or parasitoid taxa responded to tillage. While our full tillage fields tended to be smaller than our no-till fields, our comparison of models with and without field size indicates that pollinators and kleptoparasites are responding to tillage regime rather than field size. Reduced tillage may promote pollinators and kleptoparasites by destroying fewer ground-nesting bee nests (A. C. Kennedy & Schillinger, 2006; Ullmann et al., 2016), but we found intermediate tillage supported a more diverse pollinator community than no tillage. One potential explanation is that untilled soil contains a thick layer of crop stems that prevent ground nesting (Stinner & House, 1990). However, our observed impacts of tillage along with the presence of kleptoparasites suggests that many bee species nest in canola crop fields, which is often assumed to not occur (Kleijn, Rundlöf, Scheper, Smith, & Tscharntke, 2011).

Effects of tillage on natural enemies and herbivores are more well studied than for bees (Furlan, Milosavljević, Chiarini, & Benvegnù, 2021; Rowen et al., 2020; Tooker et al., 2020). Reduced tillage can benefit natural enemies by promoting weeds that provide nectar or surface mulch that provides shelter and prey (Clark, Luna, Stone, & Youngman, 1993; Stinner & House, 1990). These mechanisms are likely not operating in our system as canola provides abundant nectar, farmers controlled weeds, and crops in the region do not contribute much mulch (Hammel, 1996). When reduced tillage promotes herbivores, it mainly does so due to less soil disturbance or by promoting weeds (Rowen et al., 2020). Surface mulch can also protect aphids from predators (Hesler & Berg, 2003). None of these mechanism apply here, since we mainly sampled herbivores that reside near the tops of plants rather than in soil, farmers manged weeds, and mulch was minimal in our fields. Thus, it is not surprising that tillage did not affect natural enemies or herbivores. This highlights the necessity of studying mechanisms mediating how species’ life histories relate to food and shelter resources (Carvalheiro, Bartomeus, Rollin, Timóteo, & Tinoco, 2021).

Heterogeneous landscapes provide opportunities for consumers to exploit patchy resources (Tscharntke et al., 2012), and landscapes with more semi-natural habitat had more predators and herbivores. Predators and our main herbivores (aphids and thrips) routinely travel long distances in search of suitable habitat (Loxdale & Lushai, 1999; Schellhorn, Bianchi, & Hsu, 2014). In contrast, most pollinators are central place foragers that repeatedly return to a single nest. Indeed, bees visiting canola flowers tend to travel only a few metres from their nest (Robinson, 2019). Thus, most of the bees we collected likely were nesting in or near the canola fields we sampled, and were affected more by tillage than landscape context. This result mirrors meta-analyses that suggest that highly mobile organisms are more likely to respond to landscape-scale habitat patterns than less mobile organisms (Lichtenberg et al., 2017; Schneider et al., 2014).

While we showed that local and landscape habitat patterns affected arthropod communities, these communities minimally impacted crop yields, similar to studies that have found no relationship between multi-diversity and multifunctionality (Birkhofer et al., 2018). We did find that yield was lower in fields with higher herbivore richness and potentially in fields with fewer parasitoid taxa. Inspection of herbivore abundances at each site (Table S12) suggests two potential drivers. First, sites with higher herbivore richness could have higher pest abundance (Table S11). Second, sites with higher herbivore richness could be more likely to contain a specific damaging pest. Our data shows high-herbivore-richness fields contained large numbers of aphids, a key canola pest (Reddy, 2017). We also found more chrysomelids, curculionids, meloids, scraptiids, pentatomids, and yponomeutids in sites with higher herbivore richness. However, the only canola pests in these groups are seedpod weevils (Curculionidae), which damage a later crop stage than we sampled (Reddy, 2017). Our support for parasitoid richness increasing crop yield is mixed. Only two of three best models retained parasitoid richness, and this term was significant in only one of those models (at our chosen α = 0.05). Parasitoids can potentially enhance crop yields via pest control or even pollination as they visit flowers for nectar (Benelli et al., 2017). Ecosystem service provisioning by parasitoids is less well understood than benefits of predators and pollinators or herbivore damage (Holland et al., 2017; Noriega et al., 2018).

Abundance and diversity of pollinators and predators also did not affect yield. Canola has high variability in pollinator dependence (Ouvrard & Jacquemart, 2019), and the varieties in our study may not be pollinator dependent (Perrot, Gaba, Roncoroni, Gautier, & Bretagnolle, 2018), or the study region may be windy enough to ensure pollen dispersal. Variation in pollinator dependence could explain differences between our results and studies that find increased yield of oilseed crops with higher pollinator abundance (Catarino, Bretagnolle, Perrot, Vialloux, & Gaba, 2019). Another potential explanation is that we measured yield across the entire field but insects only along the field perimeter (due to logistics constraints). Benefits from pollinators and predators may have also been limited by insecticide use (neonicotinoid treated seeds) that may control pests and may reduce pollinator abundance. It is also possible that pollinators or predators correlate with common measures of single ecosystem services, such as pollen deposition or consumption of sentinel pests on a small subset of plants. If such patterns were present, they did not scale up to the entire field.

Tillage regime did affect crop yields, similar to studies showing reduced tillage reducing yield for oilseed rape and other crops (Lundin, 2019; Tamburini et al., 2020). This impact was direct, and not indirect via changes to the pollinator community. Indeed, multiple sustainability-oriented farming practices sometimes result in lower yield than their conventional counterparts (O. M. Smith et al., 2020; Tamburini et al., 2020). Despite yield loss seen here, reduced tillage can provide other benefits such as improving soil infiltration, reducing erosion, decreasing evaporative water loss from soil, and improving soil quality (Hammel, 1996; A. C. Kennedy & Schillinger, 2006). Reducing tillage can also reduce fuel, labour, and machine maintenance costs (although it does require up-front investment in purchasing and calibrating specialized seeding equipment, and sometimes increased fertilizing; Brown et al., 2009). These factors might ultimately increase yield of other crops or reduce farmers’ costs. This highlights the complex decisions that underlie farm management.

Overall, our study highlights the need to understand how biodiversity patterns and crop yields are simultaneously affected by multiple mechanisms, including via soil management, at various scales. We showed that tillage impacted pollinators, while landscape context strongly affected predators and herbivores. These differences likely reflect natural history differences among functional groups. However, these habitat impacts on biodiversity minimally impacted yield. It is often assumed that enhancing biodiversity promotes ecosystem services, although evidence from arthropod-mediated ecosystem services such as pollination and pest control is mixed (e.g., Birkhofer et al., 2018; Dainese et al., 2019; Ricketts et al., 2016). Without clear evidence that a conservation action such as reduced tillage is likely to increase crop yield, adoption by farmers may remain low (Kleijn et al., 2019). Thus, data-driven management of agricultural landscapes to simultaneously support natural biodiversity and ecosystem services and boost crop yield requires much more research to determine the contexts in which given management practices, and soil diversification practices in particular, do or do not meet this multi-faceted goal.

## Author Contribution

EML and DWC conceived research. EML conducted experiments. EML and IM identified arthropods. EML analysed data and conducted statistical analyses. EML, AJC and DWC wrote the manuscript. DWC secured funding. All authors read and approved the final manuscript.

## Data Availability Statement

Data and code are available via Zenodo at DOI 10.5281/zenodo.8253578.

## Acknowledgements

We thank K. Fillion, L. Rafferty, G. Smetzler, and D. Elicious for field and lab work, the growers who provided access to field sites, and three anonymous reviewers. Funding was provided by the Western Oilseed Cropping Systems program and USDA Hatch grant 1014754.

