## Supplementary material for "Differential effects of soil conservation practices on arthropods and crop yields": Table S

**Table S1:** Information about the sites we sampled.

| **Site code** | **Approximate latitude** | **Approximate longitude** | **Tillage regime** | **Year sampled** | **Dates sampled** |
|---|---|---|---|---|---|
| Cl1 | 46.8 | -117.2 | Intermediate | 2013 | 6/28-29 |
| Cl2 | 46.5 | -117.1 | Intermediate | 2013 | 7/3-4 |
| Co1 | 46.8 | -117.4 | Intermediate | 2013 | 6/30-7/1 |
| Dr1 | 46.5 | -117.1 | No | 2013 | 7/2-3 |
| Hi1 | 46.8 | -117.3 | Intermediate | 2013 | 6/17-18 |
| Hi2 | 46.8 | -117.3 | Intermediate | 2013 | 7/8-9 |
| Je1 | 46.6 | -117.0 | No | 2013 | 7/4-5 |
| Je2 | 46.6 | -117.0 | No | 2013 | 6/22-23 |
| Od1 | 46.6 | -116.9 | No | 2013 | 7/5-6 |
| Pc2 | 46.8 | -117.2 | Intermediate | 2013 | 6/27-28 |
| St3 | 46.8 | -117.8 | No | 2013 | 6/13-14 |
| St7 | 46.8 | -117.8 | No | 2013 | 6/12-13 |
| UK | 46.6 | -117.0 | Full | 2013 | 7/6-7 |
| UP | 46.7 | -117.0 | Full | 2013 | 7/7-8 |
| Wi1 | 46.7 | -117.3 | Full | 2013 | 7/1-2 |
| Be2 | 46.5 | -116.9 | Intermediate | 2014 | 6/23-24 |
| Cl4 | 46.8 | -117.2 | Intermediate | 2014 | 7/3-4 |
| Co4 | 46.8 | -117.4 | Intermediate | 2014 | 6/20-21 |
| Co5 | 46.8 | -117.4 | Intermediate | 2014 | 6/19-20 |
| Dr3 | 46.6 | -117.1 | Full | 2014 | 7/1-2 |
| Je3 | 46.6 | -117.0 | Intermediate | 2014 | 6/30-7/1 |
| Je4 | 46.6 | -117.0 | Intermediate | 2014 | 6/24-25 |
| Od2 | 46.5 | -116.8 | No | 2014 | 6/24-25 |
| Ol2 | 47.1 | -117.1 | No | 2014 | 7/9-10 |
| Ol3 | 46.8 | -117.8 | No | 2014 | 6/11-12 |
| Pc4 | 46.8 | -117.2 | Intermediate | 2014 | 7/2-3 |
| St8 | 46.8 | -117.7 | No | 2014 | 6/18-19 |
| St10 | 46.8 | -117.8 | No | 2014 | 6/18-19 |
| UK2 | 46.6 | -117.0 | Full | 2014 | 7/8-9 |
| UP4 | 46.7 | -117.0 | Full | 2014 | 7/7-8 |

**Table S2:** Taxonomic groups we sampled, their functional group categorization, and how many morphotaxa within that group are in our dataset. A $^{P}$ indicates a group known to be a major canola pest.

| **Taxon** | **Functional group** | **# morphotaxa** | **# individuals** |
|---|---|---|---|
| Coleoptera Cerambycidae | Herbivore | 1 | 68 |
| Coleoptera Chrysomelidae | Herbivore$^{P}$ | 1 | 576 |
| Coleoptera Curculionidae | Herbivore$^{P}$ | 1 | 128 |
| Coleoptera Dermestidae | Herbivore | 1 | 1 |
| Coleoptera Meloidae | Herbivore | 1 | 74 |
| Coleoptera Scarabaeidae | Herbivore | 1 | 1 |
| Coleoptera Scraptiidae | Herbivore | 1 | 54 |
| Diptera Sciaroidea | Herbivore | 1 | 2,041 |
| Hemiptera Aphididae | Herbivore$^{P}$ | 1 | 4,078 |
| Hemiptera Berytidae | Herbivore | 1 | 1 |
| Hemiptera Cercopidae | Herbivore | 1 | 1 |
| Hemiptera Cicadellidae | Herbivore | 1 | 182 |
| Hemiptera Coreoidea | Herbivore | 1 | 1 |
| Hemiptera Miridae | Herbivore | 1 | 248 |
| Hemiptera Miridae *Lygus* | Herbivore$^{P}$ | 1 | 477 |
| Hemiptera Pentatomidae | Herbivore | 1 | 5 |
| Lepidoptera Yponomeutoidea caterpillar | Herbivore | 1 | 130 |
| Orthoptera Acrididae | Herbivore | 1 | 5 |
| Orthoptera Gryllidae | Herbivore | 1 | 1 |
| Thysanoptera Thripidae | Herbivore$^{P}$ | 1 | 7,184 |
| Hymenoptera Apidae *Nomada* | Kleptoparasite | 3 | 91 |
| Hymenoptera Chrysididae | Kleptoparasite | 1 | 8 |
| Hymenoptera Crabronidae Nyssonini | Kleptoparasite | 2 | 24 |
| Hymenoptera Halictidae *Sphecodes* | Kleptoparasite | 2 | 31 |
| Diptera Conopidae | Parasitoid | 1 | 217 |
| Hymenoptera Braconidae | Parasitoid | 1 | 121 |
| Hymenoptera Chalcidoidea | Parasitoid | 1 | 727 |
| Hymenoptera Ichneumonidae | Parasitoid | 1 | 7 |
| Hymenoptera Pompilidae | Parasitoid | 1 | 6 |
| Hymenoptera Sapygidae *Sapyga pumila* | Parasitoid | 1 | 2 |
| Diptera Acroceridae | Pollinator | 1 | 3 |
| Diptera Bombyliidae | Pollinator | 1 | 5 |

| **Taxon** | **Functional group** | **# morphotaxa** | **# individuals** |
|---|---|---|---|
| Diptera Syrphidae | Pollinator | 2 | 9 |
| Hymenoptera Andrenidae | Pollinator | 2 | 245 |
| Hymenoptera Apidae Eucerini | Pollinator | 11 | 624 |
| Hymenoptera Apidae | Pollinator | 3 | 456 |
| Hymenoptera Apidae *Ceratina* | Pollinator | 2 | 28 |
| Hymenoptera Apidae *Apis mellifera* | Pollinator | 1 | 39 |
| Hymenoptera Apidae *Bombus appositus* | Pollinator | 1 | 6 |
| Hymenoptera Apidae *Bombus bifarius* | Pollinator | 1 | 19 |
| Hymenoptera Apidae *Bombus centralis* | Pollinator | 1 | 16 |
| Hymenoptera Apidae *Bombus fernaldae* | Pollinator | 1 | 1 |
| Hymenoptera Apidae *Bombus fervidus* | Pollinator | 1 | 16 |
| Hymenoptera Apidae *Bombus griseocolis* | Pollinator | 1 | 10 |
| Hymenoptera Apidae *Bombus insularis* | Pollinator | 1 | 1 |
| Hymenoptera Apidae *Bombus mixtus* | Pollinator | 1 | 5 |
| Hymenoptera Apidae *Bombus nevadensis* | Pollinator | 1 | 165 |
| Hymenoptera Apidae *Bombus occidentalis* | Pollinator | 1 | 6 |
| Hymenoptera Apidae *Bombus pensylvanicus* | Pollinator | 1 | 2 |
| Hymenoptera Apidae *Bombus rufocinctus* | Pollinator | 1 | 20 |
| Hymenoptera Apidae *Bombus vagans* | Pollinator | 1 | 1 |
| Hymenoptera Colletidae | Pollinator | 2 | 16 |
| Hymenoptera Halictidae *Agapostemon* | Pollinator | 1 | 581 |
| Hymenoptera Halictidae *Agapostemon virescens* | Pollinator | 1 | 264 |
| Hymenoptera Halictidae *Lasioglossum* | Pollinator | 4 | 836 |
| Hymenoptera Halictidae Halictini | Pollinator | 3 | 334 |
| Hymenoptera Megachilidae | Pollinator | 10 | 243 |
| Lepidoptera.Hesperiidae | Pollinator | 1 | 1 |
| Lepidoptera Noctuidae | Pollinator | 1 | 18 |
| Lepidoptera Nymphalidae | Pollinator | 1 | 6 |
| Lepidoptera Pieridae | Pollinator | 1 | 208 |
| Lepidoptera Pyralidae | Pollinator | 1 | 1 |
| Lepidoptera Sphingidae | Pollinator | 1 | 4 |

| **Taxon** | **Functional group** | **# morphotaxa** | **# individuals** |
|---|---|---|---|
| Lepidoptera Tortricidae | Pollinator | 1 | 1 |
| Lepidoptera Yponomeutoidea adult | Pollinator | 1 | 26 |
| Araneae Araneomorphae | Predator | 1 | 80 |
| Coleoptera Carabidae | Predator | 1 | 7 |
| Coleoptera Cleridae | Predator | 1 | 6 |
| Coleoptera Coccinellidae | Predator | 1 | 54 |
| Coleoptera Coccinellidae *Harmonia* | Predator | 1 | 64 |
| Coleoptera Melyridae | Predator | 1 | 229 |
| Coleoptera Staphylinidae | Predator | 1 | 7 |
| Dermaptera Forficulidae | Predator | 1 | 5 |
| Hemiptera.Anthocoridae | Predator | 1 | 47 |
| Hemiptera.Geocoridae | Predator | 1 | 33 |
| Hemiptera Nabidae *Nabis* | Predator | 1 | 1 |
| Hemiptera.Reduviidae | Predator | 1 | 4 |
| Hymenoptera Crabronidae Crabronini | Predator | 2 | 4 |
| Hymenoptera Crabronidae | Predator | 2 | 14 |
| Hymenoptera Crabronidae Larrini | Predator | 2 | 9 |
| Hymenoptera Crabronidae Cercerini | Predator | 1 | 5 |
| Hymenoptera Crabronidae Philanthini | Predator | 1 | 1 |
| Hymenoptera Formicidae | Predator | 1 | 24 |
| Hymenoptera Sphecidae | Predator | 1 | 102 |
| Hymenoptera Sphecidae *Pemphredon* | Predator | 1 | 1 |
| Hymenoptera Vespidae | Predator | 6 | 24 |
| Neuroptera.Chrysopidae | Predator | 1 | 15 |
| Neuroptera Chrysopidae *Chrysopa* | Predator | 1 | 2 |
| Odonata Coenagrionidae | Predator | 1 | 1 |

**Table S3:** Variance inflation factors (VIFs) for each full model that was the basis for model selection. We assessed collinearity using models that excluded the interaction between tillage and the proportion of semi-natural habitat within 1km of the sampling location.

| Functional group | Community metric | Tillage VIF | Proportion semi-natural habitat VIF | Year VIF | Degree days before sampling VIF | Cumulative precipitation before sampling VIF | Field size (ha planted) VIF |
|---|---|---|---|---|---|---|---|
| Herbivore | Abundance | 1.94 | 1.40 | 1.93 | 1.47 | 1.97 | 1.99 |
| Kleptoparasite | Abundance | 1.71 | 1.68 | 1.88 | 1.41 | 2.24 | 1.97 |
| Parasitoid | Abundance | 2.01 | 1.42 | 1.91 | 1.48 | 1.97 | 2.00 |
| Pollinator | Abundance | 1.93 | 1.41 | 1.93 | 1.47 | 1.99 | 1.99 |
| Predator | Abundance | 2.12 | 1.48 | 1.96 | 1.49 | 2.07 | 2.05 |
| Natural enemy | Abundance | 1.98 | 1.42 | 1.92 | 1.47 | 1.99 | 2.00 |
| Herbivore | Richness | 1.93 | 1.40 | 1.93 | 1.47 | 1.97 | 1.99 |
| Kleptoparasite | Richness | 1.93 | 1.40 | 1.93 | 1.47 | 1.97 | 1.99 |
| Parasitoid | Richness | 1.93 | 1.40 | 1.93 | 1.47 | 1.97 | 1.99 |
| Pollinator | Richness | 1.93 | 1.40 | 1.93 | 1.47 | 1.97 | 1.99 |
| Predator | Richness | 1.93 | 1.40 | 1.93 | 1.47 | 1.97 | 1.99 |
| Natural enemy | Richness | 1.93 | 1.40 | 1.93 | 1.47 | 1.97 | 1.99 |
| Herbivore | Evenness | 1.93 | 1.40 | 1.93 | 1.47 | 1.97 | 1.99 |
| Pollinator | Evenness | 1.93 | 1.40 | 1.93 | 1.47 | 1.97 | 1.99 |
| Predator | Evenness | 2.24 | 1.46 | 2.17 | 1.45 | 2.33 | 2.09 |
| Natural enemy | Evenness | 1.93 | 1.40 | 1.93 | 1.47 | 1.97 | 1.99 |

**Table S4:** Coefficients and test statistics for best-fit models investigating effects of local and landscape habitat on insect abundance. ΔAICc indicates the difference between a given model's AICc value and that of the model with the smallest AICc. Shading separates models.

| **Functional group** | **Model term** | **Coefficient** | **Coefficient standard error** | **Likelihood ratio $\chi^2$** | **df** | ***P*-value** | **ΔAICc** |
|---|---|---|---|---|---|---|---|
| Herbivore – best model 1 | Intercept | 4.92 | 0.17 | | | | 0 |
| Herbivore – best model 1 | Proportion semi-natural habitat | 4.51 | 1.87 | 6.67 | 1 | 0.01 | |
| Herbivore – best model 1 | Year | 1.46 | 0.26 | 29.42 | 1 | <0.0001 | |
| Kleptoparasite – best model 1 | Intercept | 5.13 | 1.24 | | | | 0 |
| Kleptoparasite – best model 1 | Year | 1.13 | 0.31 | 14.32 | 1 | 0.0002 | |
| Kleptoparasite – best model 1 | Degree days before sampling | -0.01 | 0.002 | 3.72 | 1 | 0.054 | |
| Kleptoparasite – best model 1 | Cumulative precipitation before sampling | -0.01 | 0.003 | 9.27 | 1 | 0.002 | |
| Kleptoparasite – best model 1 | Field size (ha planted) | 0.01 | 0.004 | 7.85 | 1 | 0.005 | |
| Kleptoparasite – best model 2 | Intercept | 2.94 | 0.70 | | | | 0.39 |
| Kleptoparasite – best model 2 | Year | 1.25 | 0.32 | 16.79 | 1 | <0.0001 | |
| Kleptoparasite – best model 2 | Cumulative precipitation before sampling | -0.01 | 0.003 | 18.79 | 1 | <0.0001 | |

| **Functional group** | **Model term** | **Coefficient** | **Coefficient standard error** | **Likelihood ratio $\chi^2$** | **df** | ***P*-value** | **ΔAICc** |
|---|---|---|---|---|---|---|---|
| Kleptoparasite – best model 2 | Field size (ha planted) | 0.01 | 0.004 | 5.18 | 1 | 0.02 | |
| Kleptoparasite – best model 3 | Intercept | 3.22 | 0.67 | | | | 0.61 |
| Kleptoparasite – best model 3 | Tillage – full till | -1.45 | 0.58 | 8.62 | 2 | 0.01 | |
| Kleptoparasite – best model 3 | Tillage – no till | 0.15 | 0.30 | | | | |
| Kleptoparasite – best model 3 | Year | 1.33 | 0.29 | 22.26 | 1 | <0.0001 | |
| Kleptoparasite – best model 3 | Cumulative precipitation before sampling | -0.01 | 0.003 | 14.05 | 1 | 0.0002 | |
| Kleptoparasite – best model 4 | Intercept | 2.80 | 0.70 | | | | 1.82 |
| Kleptoparasite – best model 4 | Tillage – full till | -1.25 | 0.59 | 5.55 | 2 | 0.06 | |
| Kleptoparasite – best model 4 | Tillage – no till | 0.08 | 0.30 | | | | |
| Kleptoparasite – best model 4 | Year | 1.22 | 0.29 | 19.11 | 1 | <0.0001 | |
| Kleptoparasite – best model 4 | Cumulative precipitation before sampling | -0.01 | 0.003 | 12.69 | 1 | 0.0004 | |
| Kleptoparasite – best model 4 | Field size (ha planted) | 0.01 | 0.004 | 2.30 | 1 | 0.13 | |
| Kleptoparasite – best constrained model 1 | Intercept | 2.80 | 0.70 | | | | 0 |

| **Functional group** | **Model term** | **Coefficient** | **Coefficient standard error** | **Likelihood ratio $\chi^2$** | **df** | ***P*-value** | **ΔAICc** |
|---|---|---|---|---|---|---|---|
| Kleptoparasite – best constrained model 1 | Tillage – full till | -1.25 | 0.59 | 5.55 | 2 | 0.06 | |
| Kleptoparasite – best constrained model 1 | Tillage – no till | 0.08 | 0.30 | | | | |
| Kleptoparasite – best constrained model 1 | Year | 1.22 | 0.29 | 19.11 | 1 | <0.0001 | |
| Kleptoparasite – best constrained model 1 | Cumulative precipitation before sampling | -0.01 | 0.003 | 12.69 | 1 | 0.0004 | |
| Kleptoparasite – best constrained model 1 | Field size (ha planted) | 0.01 | 0.004 | 2.30 | 1 | 0.13 | |
| Kleptoparasite – best constrained model 2 | Intercept | 3.84 | 0.61 | | | | 0.51 |
| Kleptoparasite – best constrained model 2 | Year | 1.47 | 0.33 | 21.70 | 1 | <0.0001 | |
| Kleptoparasite – best constrained model 2 | Cumulative precipitation before sampling | -0.01 | 0.003 | 25.09 | 1 | <0.0001 | |
| Parasitoid – best model 1 | Intercept | 6.63 | 1.27 | | | | 0 |
| Parasitoid – best model 1 | Year | 1.15 | 0.30 | 15.90 | 1 | <0.0001 | |
| Parasitoid – best model 1 | Degree days before sampling | -0.01 | 0.002 | 18.83 | 1 | <0.0001 | |

| **Functional group** | **Model term** | **Coefficient** | **Coefficient standard error** | **Likelihood ratio $\chi^2$** | **df** | ***P*-value** | **ΔAICc** |
|---|---|---|---|---|---|---|---|
| Parasitoid – best model 1 | Cumulative precipitation before sampling | 0.01 | 0.003 | 4.08 | 1 | 0.04 | |
| Parasitoid – best model 2 | Intercept | 7.20 | 1.34 | | | | 0.94 |
| Parasitoid – best model 2 | Year | 1.37 | 0.28 | 21.90 | 1 | <0.0001 | |
| Parasitoid – best model 2 | Degree days before sampling | -0.01 | 0.002 | 14.08 | 1 | 0.0002 | |
| Pollinator – best model 1 | Intercept | 6.12 | 0.44 | | | | 0 |
| Pollinator – best model 1 | Year | 0.94 | 0.21 | 18.60 | 1 | <0.0001 | |
| Pollinator – best model 1 | Cumulative precipitation before sampling | -0.01 | 0.002 | 15.85 | 1 | <0.0001 | |
| Pollinator – best model 2 | Intercept | 5.81 | 0.52 | | | | 1.58 |
| Pollinator – best model 2 | Year | 0.87 | 0.22 | 15.26 | 1 | <0.0001 | |
| Pollinator – best model 2 | Cumulative precipitation before sampling | -0.01 | 0.002 | 11.96 | 1 | 0.0006 | |
| Pollinator – best model 2 | Field size (ha planted) | 0.003 | 0.003 | 1.35 | 1 | 0.25 | |
| Predator – best model 1 | Intercept | 2.73 | 0.16 | | | | 0 |
| Predator – best model 1 | Proportion semi-natural habitat | 6.64 | 1.64 | 24.96 | 1 | <0.0001 | |

| **Functional group** | **Model term** | **Coefficient** | **Coefficient standard error** | **Likelihood ratio $\chi^2$** | **df** | ***P*-value** | **ΔAICc** |
|---|---|---|---|---|---|---|---|
| Predator – best model 1 | Year | 0.04 | 0.24 | 0.03 | 1 | 0.86 | |
| Predator – best model 2 | Intercept | 3.38 | 0.47 | | | | 0.25 |
| Predator – best model 2 | Proportion semi-natural habitat | 6.28 | 1.60 | 18.24 | 1 | <0.0001 | |
| Predator – best model 2 | Year | 0.26 | 0.25 | 0.92 | 1 | 0.34 | |
| Predator – best model 2 | Cumulative precipitation before sampling | -0.003 | 0.002 | 2.75 | 1 | 0.10 | |
| Predator – best model 3 | Intercept | 4.30 | 1.01 | | | | 0.67 |
| Predator – best model 3 | Proportion semi-natural habitat | 6.28 | 1.59 | 21.02 | 1 | <0.0001 | |
| Predator – best model 3 | Year | 0.09 | 0.23 | 0.15 | 1 | 0.70 | |
| Predator – best model 3 | Degree days before sampling | -0.003 | 0.002 | 2.31 | 1 | 0.13 | |
| Natural enemy – best model 1 | Intercept | 6.66 | 1.04 | | | | 0 |
| Natural enemy – best model 1 | Proportion semi-natural habitat | 3.13 | 1.71 | 3.64 | 1 | 0.056 | |
| Natural enemy – best model 1 | Year | 0.80 | 0.24 | 10.67 | 1 | 0.001 | |
| Natural enemy – best model 1 | Degree days before sampling | -0.01 | 0.002 | 10.51 | 1 | 0.001 | |

| **Functional group** | **Model term** | **Coefficient** | **Coefficient standard error** | **Likelihood ratio $\chi^2$** | **df** | ***P*-value** | **ΔAICc** |
|---|---|---|---|---|---|---|---|
| Natural enemy – best model 2 | Intercept | 7.14 | 1.09 | | | | 0.53 |
| Natural enemy – best model 2 | Year | 1.03 | 0.23 | 18.96 | 1 | <0.0001 | |
| Natural enemy – best model 2 | Degree days before sampling | -0.01 | 0.002 | 12.67 | 1 | 0.0004 | |

**Table S5:** Coefficients and test statistics for best-fit models investigating effects of local and landscape habitat on insect richness. Residual degrees of freedom are listed in the Intercept row for each model. ΔAICc indicates the difference between a given model's AICc value and that of the model with the smallest AICc. Shading separates models.

| **Functional group** | **Model term** | **Coefficient** | **Coefficient standard error** | **F** | **df** | ***P*-value** | **ΔAICc** |
|---|---|---|---|---|---|---|---|
| Herbivore – best model 1 | Intercept | 7.64 | 0.52 | | 27 | | 0 |
| Herbivore – best model 1 | Proportion semi-natural habitat | 9.63 | 5.83 | 2.73 | 1 | 0.11 | |
| Herbivore – best model 1 | Year | 1.05 | 0.81 | 1.71 | 1 | 0.20 | |
| Herbivore – best model 2 | Intercept | 7.73 | 0.54 | | 28 | | 0.21 |
| Herbivore – best model 2 | Year | 1.60 | 0.76 | 4.47 | 1 | 0.04 | |
| Herbivore – best model 3 | Intercept | 8.27 | 0.63 | | 27 | | 0.41 |
| Herbivore – best model 3 | Year | 1.79 | 0.75 | 5.71 | 1 | 0.02 | |
| Herbivore – best model 3 | Field size (ha planted) | -0.01 | 0.01 | 2.32 | 1 | 0.14 | |
| Herbivore – best model 4 | Intercept | 12.63 | 3.52 | | 27 | | 0.77 |
| Herbivore – best model 4 | Year | 1.56 | 0.75 | 4.37 | 1 | 0.046 | |
| Herbivore – best model 4 | Degree days before sampling | -0.01 | 0.01 | 1.98 | 1 | 0.17 | |
| Herbivore – best model 5 | Intercept | 13.11 | 3.45 | | 26 | | 1.05 |

| **Functional group** | **Model term** | **Coefficient** | **Coefficient standard error** | **F** | **df** | ***P*-value** | **ΔAICc** |
|---|---|---|---|---|---|---|---|
| Herbivore – best model 5 | Year | 1.75 | 0.74 | 5.63 | 1 | 0.03 | |
| Herbivore – best model 5 | Degree days before sampling | -0.01 | 0.01 | 2.04 | 1 | 0.17 | |
| Herbivore – best model 5 | Field size (ha planted | -0.01 | 0.01 | 2.38 | 1 | 0.14 | |
| Herbivore – best model 6 | Intercept | 11.51 | 3.36 | | 25 | | 1.10 |
| Herbivore – best model 6 | Proportion semi-natural habitat | 10.12 | 5.68 | 3.18 | 1 | 0.09 | |
| Herbivore – best model 6 | Year | 0.43 | 0.86 | 0.25 | 1 | 0.62 | |
| Herbivore – best model 6 | Degree days before sampling | -0.01 | 0.01 | 3.38 | 1 | 0.08 | |
| Herbivore – best model 6 | Cumulative precipitation before sampling | 0.01 | 0.01 | 2.84 | 1 | 0.11 | |
| Herbivore – best model 7 | Intercept | 8.09 | 0.63 | | 26 | | 1.12 |
| Herbivore – best model 7 | Proportion semi-natural habitat | 8.24 | 5.87 | 1.97 | 1 | 0.17 | |
| Herbivore – best model 7 | Year | 1.29 | 0.82 | 2.49 | 1 | 0.13 | |
| Herbivore – best model 7 | Field size (ha planted) | -0.01 | 0.01 | 1.59 | 1 | 0.22 | |
| Herbivore – best model 8 | Intercept | 11.91 | 3.47 | | 26 | | 1.17 |

| Functional group | Model term | Coefficient | Coefficient standard error | F | df | *P*-value | ΔAICc |
|---|---|---|---|---|---|---|---|
| Herbivore – best model 8 | Proportion semi-natural habitat | 8.73 | 5.81 | 2.26 | 1 | 0.15 | |
| Herbivore – best model 8 | Year | 1.07 | 0.80 | 1.79 | 1 | 0.19 | |
| Herbivore – best model 8 | Degree days before sampling | -0.01 | 0.01 | 1.54 | 1 | 0.23 | |
| Herbivore – best model 9 | Intercept | 12.39 | 3.46 | | 26 | | 1.54 |
| Herbivore – best model 9 | Year | 1.09 | 0.81 | 1.81 | 1 | 0.19 | |
| Herbivore – best model 9 | Degree days before sampling | -0.01 | 0.01 | 3.48 | 1 | 0.07 | |
| Herbivore – best model 9 | Cumulative precipitation before sampling | 0.01 | 0.01 | 1.91 | 1 | 0.18 | |
| Herbivore – best model 10 | Intercept | 6.05 | 1.66 | | 26 | | 1.75 |
| Herbivore – best model 10 | Proportion semi-natural habitat | 10.71 | 5.92 | 3.27 | 1 | 0.08 | |
| Herbivore – best model 10 | Year | 0.69 | 0.89 | 0.60 | 1 | 0.45 | |
| Herbivore – best model 10 | Cumulative precipitation before sampling | 0.01 | 0.01 | 1.01 | 1 | 0.32 | |
| Kleptoparasite – best model 1 | Intercept | 8.54 | 1.97 | | 25 | | 0 |
| Kleptoparasite – best model 1 | Year | 1.51 | 0.49 | 9.37 | 1 | 0.01 | |

| Functional group | Model term | Coefficient | Coefficient standard error | F | df | *P*-value | ΔAICc |
|---|---|---|---|---|---|---|---|
| Kleptoparasite – best model 1 | Degree days before sampling | -0.01 | 0.003 | 4.28 | 1 | 0.049 | |
| Kleptoparasite – best model 1 | Cumulative precipitation before sampling | -0.01 | 0.01 | 6.99 | 1 | 0.01 | |
| Kleptoparasite – best model 1 | Field size (ha planted) | 0.02 | 0.01 | 6.79 | 1 | 0.02 | |
| Kleptoparasite – best model 2 | Intercept | 5.16 | 1.18 | | 26 | | 1.59 |
| Kleptoparasite – best model 2 | Year | 1.78 | 0.50 | 12.49 | 1 | 0.002 | |
| Kleptoparasite – best model 2 | Cumulative precipitation before sampling | -0.02 | 0.004 | 14.23 | 1 | 0.0008 | |
| Kleptoparasite – best model 2 | Field size (ha planted) | 0.01 | 0.01 | 4.31 | 1 | 0.048 | |
| Parasitoid – best model 1 | Intercept | 2.20 | 0.23 | | 28 | | 0 |
| Parasitoid – best model 1 | Year | 0.87 | 0.32 | 7.39 | 1 | 0.01 | |
| Parasitoid – best model 2 | Intercept | 2.17 | 0.23 | | 27 | | 1.20 |
| Parasitoid – best model 2 | Proportion semi-natural habitat | 2.93 | 2.51 | 1.36 | 1 | 0.25 | |
| Parasitoid – best model 2 | Year | 0.70 | 0.35 | 4.07 | 1 | 0.054 | |
| Parasitoid – best model 3 | Intercept | 3.74 | 1.51 | | 27 | | 1.52 |

| **Functional group** | **Model term** | **Coefficient** | **Coefficient standard error** | **F** | **df** | ***P*-value** | **ΔAICc** |
|---|---|---|---|---|---|---|---|
| Parasitoid – best model 3 | Year | 0.85 | 0.32 | 7.17 | 1 | 0.01 | |
| Parasitoid – best model 3 | Degree days before sampling | -0.002 | 0.002 | 1.07 | 1 | 0.31 | |
| Pollinator – best model 1 | Intercept | 29.06 | 4.13 | | 25 | | 0 |
| Pollinator – best model 1 | Tillage – full till | -1.85 | 2.30 | 5.59 | 2 | 0.01 | |
| Pollinator – best model 1 | Tillage – no till | -6.23 | 1.87 | | | | |
| Pollinator – best model 1 | Year | 5.56 | 1.77 | 9.90 | 1 | 0.004 | |
| Pollinator – best model 1 | Cumulative precipitation before sampling | -0.03 | 0.02 | 3.42 | 1 | 0.08 | |
| Pollinator – best model 2 | Intercept | 21.95 | 1.57 | | 26 | | 0.69 |
| Pollinator – best model 2 | Tillage – full till | -3.23 | 2.28 | 4.15 | 2 | 0.03 | |
| Pollinator – best model 2 | Tillage – no till | -5.42 | 1.90 | | | | |
| Pollinator – best model 2 | Year | 4.24 | 1.69 | 6.29 | 1 | 0.02 | |
| Pollinator – best model 3 | Intercept | 20.75 | 1.77 | | 25 | | 1.66 |
| Pollinator – best model 3 | Tillage – full till | -1.99 | 2.42 | 5.12 | 2 | 0.01 | |
| Pollinator – best model 3 | Tillage – no till | -6.30 | 1.97 | | | | |

| Functional group | Model term | Coefficient | Coefficient standard error | F | df | *P*-value | ΔAICc |
|---|---|---|---|---|---|---|---|
| Pollinator – best model 3 | Year | 3.71 | 1.71 | 4.73 | 1 | 0.04 | |
| Pollinator – best model 3 | Field size (ha planted) | 0.04 | 0.03 | 1.89 | 1 | 0.18 | |
| Pollinator – best constrained model 1 | Intercept | 20.75 | 1.77 | | 25 | | 0 |
| Pollinator – best constrained model 1 | Tillage – full till | -1.99 | 2.42 | 5.12 | 2 | 0.01 | |
| Pollinator – best constrained model 1 | Tillage – no till | -6.30 | 1.97 | | | | |
| Pollinator – best constrained model 1 | Year | 3.71 | 1.71 | 4.73 | 1 | 0.04 | |
| Pollinator – best constrained model 1 | Field size (ha planted) | 0.04 | 0.03 | 1.89 | 1 | 0.18 | |
| Pollinator – best constrained model 2 | Intercept | 27.25 | 4.50 | | 24 | | 0.53 |
| Pollinator – best constrained model 2 | Tillage – full till | -1.11 | 2.42 | 6.10 | 2 | 0.007 | |
| Pollinator – best constrained model 2 | Tillage – no till | -6.78 | 1.94 | | | | |

| **Functional group** | **Model term** | **Coefficient** | **Coefficient standard error** | **F** | **df** | ***P*-value** | **ΔAICc** |
|---|---|---|---|---|---|---|---|
| Pollinator – best constrained model 2 | Year | 5.00 | 1.85 | 7.29 | 1 | 0.01 | |
| Pollinator – best constrained model 2 | Cumulative precipitation before sampling | -0.03 | 0.02 | 2.44 | 1 | 0.13 | |
| Pollinator – best constrained model 2 | Field size (ha planted) | 0.03 | 0.03 | 1.02 | 1 | 0.32 | |
| Pollinator – best constrained model 3 | Intercept | 19.13 | 1.32 | | 28 | | 1.77 |
| Pollinator – best constrained model 3 | Year | 4.60 | 1.87 | 6.08 | 1 | 0.02 | |
| Predator – best model 1 | Intercept | 15.24 | 3.77 | | 27 | | 0 |
| Predator – best model 1 | Year | 0.92 | 0.80 | 1.32 | 1 | 0.26 | |
| Predator – best model 1 | Degree days before sampling | -0.01 | 0.01 | 6.04 | 1 | 0.02 | |
| Predator – best model 2 | Intercept | 15.49 | 3.73 | | 26 | | 0.89 |
| Predator – best model 2 | Year | 1.41 | 0.87 | 2.64 | 1 | 0.12 | |
| Predator – best model 2 | Degree days before sampling | -0.01 | 0.01 | 3.04 | 1 | 0.09 | |
| Predator – best model 2 | Cumulative precipitation before sampling | -0.01 | 0.01 | 1.80 | 1 | 0.19 | |

| **Functional group** | **Model term** | **Coefficient** | **Coefficient standard error** | **F** | **df** | ***P*-value** | **ΔAICc** |
|---|---|---|---|---|---|---|---|
| Predator – best model 3 | Intercept | 9.74 | 1.80 | | 27 | | 1.31 |
| Predator – best model 3 | Year | 1.72 | 0.88 | 3.80 | 1 | 0.06 | |
| Predator – best model 3 | Cumulative precipitation before sampling | -0.02 | 0.01 | 4.63 | 1 | 0.04 | |
| Predator – best model 4 | Intercept | 14.76 | 3.82 | | 26 | | 1.95 |
| Predator – best model 4 | Proportion semi-natural habitat | 5.85 | 6.40 | 0.83 | 1 | 0.37 | |
| Predator – best model 4 | Year | 0.59 | 0.88 | 0.45 | 1 | 0.51 | |
| Predator – best model 4 | Degree days before sampling | -0.01 | 0.01 | 5.38 | 1 | 0.03 | |
| Natural enemy – best model 1 | Intercept | 18.98 | 4.30 | | 27 | | 0 |
| Natural enemy – best model 1 | Year | 1.77 | 0.91 | 3.79 | 1 | 0.06 | |
| Natural enemy – best model 1 | Degree days before sampling | -0.02 | 0.01 | 6.35 | 1 | 0.02 | |
| Natural enemy – best model 2 | Intercept | 18.27 | 4.31 | | 26 | | 1.34 |
| Natural enemy – best model 2 | Proportion semi-natural habitat | 8.50 | 7.21 | 1.39 | 1 | 0.25 | |
| Natural enemy – best model 2 | Year | 1.30 | 0.99 | 1.71 | 1 | 0.20 | |

| **Functional group** | **Model term** | **Coefficient** | **Coefficient standard error** | **F** | **df** | ***P*-value** | **ΔAICc** |
|---|---|---|---|---|---|---|---|
| Natural enemy – best model 2 | Degree days before sampling | -0.02 | 0.01 | 5.63 | 1 | 0.03 | |
| Natural enemy – best model 3 | Intercept | 19.21 | 4.29 | | 26 | | 1.56 |
| Natural enemy – best model 3 | Year | 2.23 | 1.00 | 4.98 | 1 | 0.04 | |
| Natural enemy – best model 3 | Degree days before sampling | -0.01 | 0.01 | 3.52 | 1 | 0.07 | |
| Natural enemy – best model 3 | Cumulative precipitation before sampling | -0.01 | 0.01 | 1.19 | 1 | 0.29 | |

**Table S6:** Coefficients and test statistics for best-fit models investigating effects of local and landscape habitat on insect evenness. Herbivore evenness was square root transformed. Residual degrees of freedom are listed in the Intercept row for each model. ΔAICc indicates the difference between a given model's AICc value and that of the model with the smallest AICc. Shading separates models.

| Functional group | Model term | Coefficient | Coefficient standard error | F | df | *P*-value | ΔAICc |
|---|---|---|---|---|---|---|---|
| Herbivore – best model | Intercept | 0.60 | 0.03 | | 28 | | 0 |
| Herbivore – best model | Year | -0.17 | 0.04 | 0.20 | 1 | 0.0002 | |
| Pollinator – best model 1 | Intercept | 0.34 | 0.09 | | 27 | | 0 |
| Pollinator – best model 1 | Year | -0.12 | 0.05 | 6.68 | 1 | 0.02 | |
| Pollinator – best model 1 | Cumulative precipitation before sampling | 0.001 | 0.0004 | 4.28 | 1 | 0.048 | |
| Pollinator – best model 2 | Intercept | 0.58 | 0.04 | | 26 | | 0.35 |
| Pollinator – best model 2 | Proportion semi-natural habitat | -0.60 | 0.34 | 3.22 | 1 | 0.08 | |
| Pollinator – best model 2 | Year | -0.03 | 0.05 | 0.49 | 1 | 0.49 | |
| Pollinator – best model 2 | Field size (planted ha) | -0.001 | 0.001 | 4.80 | 1 | 0.04 | |
| Pollinator – best model 3 | Intercept | 0.56 | 0.03 | | 27 | | 0.96 |
| Pollinator – best model 3 | Year | -0.07 | 0.04 | 2.50 | 1 | 0.13 | |
| Pollinator – best model 3 | Field size (ha planted) | -0.001 | 0.001 | 3.30 | 1 | 0.08 | |

| Functional group | Model term | Coefficient | Coefficient standard error | F | df | *P*-value | ΔAICc |
|---|---|---|---|---|---|---|---|
| Pollinator – best model 4 | Intercept | 0.41 | 0.11 | | 26 | | 1.64 |
| Pollinator – best model 4 | Year | -0.10 | 0.05 | 4.34 | 1 | 0.047 | |
| Pollinator – best model 4 | Cumulative precipitation before sampling | 0.001 | 0.0004 | 2.00 | 1 | 0.17 | |
| Pollinator – best model 4 | Field size (ha planted) | -0.001 | 0.001 | 1.12 | 1 | 0.30 | |
| Pollinator – best model 5 | Intercept | 0.36 | 0.10 | | 26 | | 1.73 |
| Pollinator – best model 5 | Proportion semi-natural habitat | -0.35 | 0.34 | 1.04 | 1 | 0.32 | |
| Pollinator – best model 5 | Year | -0.10 | 0.05 | 3.49 | 1 | 0.07 | |
| Pollinator – best model 5 | Cumulative precipitation before sampling | 0.001 | 0.0004 | 3.43 | 1 | 0.08 | |
| Pollinator – best model 6 | Intercept | 0.53 | 0.03 | | 28 | | 1.74 |
| Pollinator – best model 6 | Year | -0.08 | 0.05 | 3.40 | 1 | 0.08 | |
| Predator – best model | Intercept | 0.74 | 0.04 | | 25 | | 0 |
| Predator – best model | Proportion semi-natural habitat | -1.28 | 0.45 | 8.11 | 1 | 0.009 | |
| Predator – best model | Year | -0.02 | 0.06 | 0.08 | 1 | 0.78 | |

| **Functional group** | **Model term** | **Coefficient** | **Coefficient standard error** | **F** | **df** | ***P*-value** | **ΔAICc** |
|---|---|---|---|---|---|---|---|
| Natural enemy – best model 1 | Intercept | 0.65 | 0.04 | | 28 | | 0 |
| Natural enemy – best model 1 | Year | -0.22 | 0.06 | 13.88 | 1 | 0.0009 | |
| Natural enemy – best model 2 | Intercept | 0.66 | 0.04 | | 27 | | 0.38 |
| Natural enemy – best model 2 | Proportion semi-natural habitat | -0.69 | 0.47 | 2.15 | 1 | 0.15 | |
| Natural enemy – best model 2 | Year | -0.19 | 0.07 | 8.21 | 1 | 0.008 | |
| Natural enemy – best model 3 | Intercept | 0.40 | 0.29 | | 27 | | 1.85 |
| Natural enemy – best model 3 | Year | -0.22 | 0.06 | 13.47 | 1 | 0.001 | |
| Natural enemy – best model 3 | Degree days before sampling | 0.0004 | 0.0004 | 0.76 | 1 | 0.39 | |

**Table S7:** Variance inflation factors (VIFs) for each full yield model that was the basis for model selection. We assessed collinearity using models that excluded the interaction between tillage and the proportion of semi-natural habitat within 1km of the sampling location. Evenness regressions did not include kleptoparasites or parasitoids, which had too few sites with sufficient richness to calculate evenness. Because some variables in full models had moderate VIFs (between 5 and 10, two common VIF thresholds; James et al. 2022), we also show VIFs for the best models.

| **Community metric** | **Tillage VIF** | **Proportion semi-natural habitat VIF** | **Year VIF** | **Field size VIF** | **Herbivore VIF** | **Kleptoparasite VIF** | **Parasitoid VIF** | **Pollinator VIF** | **Predator VIF** |
|---|---|---|---|---|---|---|---|---|---|
| Abundance – full model | 1.74 | 7.38 | 2.45 | NA | 5.73 | 3.20 | 1.78 | 2.92 | 6.33 |
| Abundance – best model 1 | 1.01 | NA | 1.01 | NA | NA | NA | NA | NA | NA |
| Abundance – best model 2 | 1.04 | NA | 1.30 | NA | NA | NA | 1.34 | NA | NA |
| Abundance – best model 3 | 1.04 | NA | 1.10 | NA | NA | NA | NA | NA | 1.12 |
| Abundance – constrained full model | 2.20 | 7.38 | 2.54 | 1.91 | 5.79 | 3.32 | 1.79 | 2.98 | 6.46 |
| Abundance – constrained best model 1 | 1.46 | NA | 1.06 | 1.50 | NA | NA | NA | NA | NA |
| Abundance – constrained best model 1 | 1.53 | NA | 1.39 | 1.52 | NA | NA | 1.36 | NA | NA |
| Richness – full model | 3.12 | 1.41 | 2.05 | NA | 2.79 | 4.19 | 1.54 | 2.72 | 3.35 |
| Richness – best model 1 | 1.07 | NA | 1.36 | NA | 1.23 | NA | 1.40 | NA | NA |
| Richness – best model 2 | 1.41 | NA | 1.62 | NA | 1.35 | NA | 1.50 | NA | 1.84 |

| **Community metric** | **Tillage VIF** | **Proportion semi-natural habitat VIF** | **Year VIF** | **Field size VIF** | **Herbivore VIF** | **Kleptoparasite VIF** | **Parasitoid VIF** | **Pollinator VIF** | **Predator VIF** |
|---|---|---|---|---|---|---|---|---|---|
| Richness – best model 3 | 1.01 | NA | 1.17 | NA | 1.17 | NA | NA | NA | NA |
| Richness – constrained full model | 3.47 | 1.52 | 2.11 | 1.99 | 2.83 | 4.50 | 1.55 | 2.73 | 3.35 |
| Richness – constrained best model 1 | 1.55 | NA | 1.48 | 1.64 | 1.35 | NA | 1.40 | NA | NA |
| Richness – best model 2 | 1.48 | NA | 1.30 | 1.64 | 1.28 | NA | NA | NA | NA |
| Richness – best model 3 | 2.14 | NA | 1.68 | 1.71 | 1.43 | NA | 1.50 | 1.92 | NA |
| Evenness – full model | 1.56 | 1.71 | 2.10 | NA | 1.86 | NA | NA | 1.42 | 1.74 |
| Evenness – best model | 1.01 | NA | 1.01 | NA | NA | NA | NA | NA | NA |
| Evenness – constrained full model | 2.31 | 2.10 | 2.33 | 1.96 | 1.89 | NA | NA | 1.53 | 1.86 |
| Evenness – constrained best model | 1.42 | NA | 1.08 | 1.48 | NA | NA | NA | NA | NA |

**Table S8:** Coefficients and test statistics for best-fit models investigating effects of local and landscape habitat, and arthropod communities, on crop yield (kg/ha). Residual degrees of freedom are listed in the Intercept row for each model. ΔAICc indicates the difference between a given model's AICc value and that of the model with the smallest AICc. Shading separates models.

| **Model** | **Model term** | **Coefficient** | **Coefficient standard error** | **F** | **df** | ***P*-value** | **ΔAICc** |
|---|---|---|---|---|---|---|---|
| Abundance – best 1 | Intercept | 2477.6 | 177.3 | | 26 | | 0 |
| Abundance – best 1 | Tillage – full till | 260.2 | 257.3 | 6.54 | 2 | 0.005 | |
| Abundance – best 1 | Tillage – no till | -604.9 | 214.1 | | | | |
| Abundance – best 1 | Year | -410.2 | 190.8 | 4.62 | 1 | 0.04 | |
| Abundance – best 2 | Intercept | 2404.10 | 186.81 | | 25 | | 1.54 |
| Abundance – best 2 | Tillage – full till | 316.26 | 259.85 | 6.88 | 2 | 0.004 | |
| Abundance – best 2 | Tillage – no till | -588.39 | 213.00 | | | | |
| Abundance – best 2 | Year | -530.77 | 215.43 | 6.07 | 1 | 0.02 | |
| Abundance – best 2 | Parasitoid abundance | 3.27 | 2.75 | 1.38 | 1 | 0.25 | |
| Abundance – best 3 | Intercept | 2509.37 | 179.73 | | 25 | | 1.89 |
| Abundance – best 3 | Tillage – full till | 283.06 | 257.90 | 5.92 | 2 | 0.008 | |
| Abundance – best 3 | Tillage – no till | -561.28 | 217.90 | | | | |
| Abundance – best 3 | Year | -350.59 | 199.07 | 3.10 | 1 | 0.09 | |

| Model | Model term | Coefficient | Coefficient standard error | F | df | *P*-value | ΔAICc |
|---|---|---|---|---|---|---|---|
| Abundance – best 3 | Predator abundance | -3.34 | 3.22 | 1.07 | 1 | 0.31 | |
| Abundance – constrained best 1 | Intercept | 2439.98 | 207.18 | | 25 | | 0 |
| Abundance – constrained best 1 | Tillage – full till | 299.21 | 282.31 | 5.19 | 2 | 0.01 | |
| Abundance – constrained best 1 | Tillage – no till | -632.45 | 230.21 | | | | |
| Abundance – constrained best 1 | Year | -426.74 | 199.19 | 4.59 | 1 | 0.04 | |
| Abundance – constrained best 1 | Field size (planted ha) | 1.08 | 2.94 | 0.14 | 1 | 0.72 | |
| Abundance – constrained best 2 | Intercept | 2347.26 | 219.03 | | 24 | | 1.65 |
| Abundance – constrained best 2 | Tillage – full till | 374.18 | 286.43 | 5.64 | 2 | 0.01 | |
| Abundance – constrained best 2 | Tillage – no till | -626.16 | 228.13 | | | | |
| Abundance – constrained best 2 | Year | -560.68 | 226.13 | 6.15 | 1 | 0.02 | |

| Model | Model term | Coefficient | Coefficient standard error | F | df | *P*-value | ΔAICc |
|---|---|---|---|---|---|---|---|
| Abundance – constrained best 2 | Field size (planted ha) | 1.52 | 2.94 | 0.27 | 1 | 0.61 | |
| Abundance – constrained best 2 | Parasitoid abundance | 3.42 | 2.82 | 1.47 | 1 | 0.24 | |
| Richness – best 1 | Intercept | 2910.11 | 396.65 | | 24 | | 0 |
| Richness – best 1 | Tillage – full till | 402.27 | 232.21 | 9.44 | 2 | 0.001 | |
| Richness – best 1 | Tillage – no till | -562.45 | 189.10 | | | | |
| Richness – best 1 | Year | -393.48 | 194.33 | 4.10 | 1 | 0.054 | |
| Richness – best 1 | Herbivore richness | -120.56 | 42.86 | 7.91 | 1 | 0.01 | |
| Richness – best 1 | Parasitoid richness | 206.55 | 103.91 | 3.95 | 1 | 0.058 | |
| Richness – best 2 | Intercept | 3626.38 | 614.56 | | 23 | | 0.97 |
| Richness – best 2 | Tillage – full till | 331.42 | 231.27 | 11.02 | 2 | 0.0004 | |
| Richness – best 2 | Tillage – no till | -718.07 | 211.57 | | | | |
| Richness – best 2 | Year | -269.40 | 206.74 | 1.70 | 1 | 0.21 | |
| Richness – best 2 | Herbivore richness | -140.24 | 43.80 | 10.25 | 1 | 0.004 | |
| Richness – best 2 | Parasitoid richness | 248.82 | 105.16 | 5.60 | 1 | 0.03 | |
| Richness – best 2 | Pollinator richness | -30.35 | 20.24 | 2.25 | 1 | 0.15 | |
| Richness – best 3 | Intercept | 3256.26 | 376.83 | | 25 | | 1.13 |

| **Model** | **Model term** | **Coefficient** | **Coefficient standard error** | **F** | **df** | ***P*-value** | **ΔAICc** |
|---|---|---|---|---|---|---|---|
| Richness – best 3 | Tillage – full till | 296.29 | 238.97 | 8.09 | 2 | 0.002 | |
| Richness – best 3 | Tillage – no till | -609.02 | 198.41 | | | | |
| Richness – best 3 | Year | -248.22 | 190.39 | 1.70 | 1 | 0.20 | |
| Richness – best 3 | Herbivore richness | -101.41 | 44.16 | 5.27 | 1 | 0.03 | |
| Richness – constrained best 1 | Intercept | 2981.59 | 444.60 | | 23 | | 0 |
| Richness – constrained best 1 | Tillage – full till | 367.61 | 253.05 | 5.64 | 2 | 0.01 | |
| Richness – constrained best 1 | Tillage – no till | -535.81 | 204.64 | | | | |
| Richness – constrained best 1 | Year | -370.93 | 206.38 | 3.23 | 1 | 0.09 | |
| Richness – constrained best 1 | Field size (planted ha) | -1.04 | 2.70 | 0.15 | 1 | 0.70 | |
| Richness – constrained best 1 | Herbivore richness | -125.77 | 45.70 | 7.68 | 1 | 0.01 | |
| Richness – constrained best 1 | Parasitoid richness | 208.45 | 105.92 | 387 | 1 | 0.06 | |

| **Model** | **Model term** | **Coefficient** | **Coefficient standard error** | **F** | **df** | ***P*-value** | **ΔAICc** |
|---|---|---|---|---|---|---|---|
| Richness – constrained best 2 | Intercept | 3313.09 | 435.38 | | 24 | | 0.90 |
| Richness – constrained best 2 | Tillage – full till | 269.16 | 262.48 | 5.09 | 2 | 0.01 | |
| Richness – constrained best 2 | Tillage – no till | -589.07 | 214.63 | | | | |
| Richness – constrained best 2 | Year | -230.03 | 204.82 | 1.26 | 1 | 0.27 | |
| Richness – constrained best 2 | Field size (planted ha) | -0.79 | 2.86 | 0.08 | 1 | 0.78 | |
| Richness – constrained best 2 | Herbivore richness | -105.24 | 47.08 | 5.00 | 1 | 0.04 | |
| Richness – constrained best 3 | Intercept | 3634.32 | 633.81 | | 22 | | 1.52 |
| Richness – constrained best 3 | Tillage – full till | 323.83 | 249.60 | 6.86 | 2 | 0.005 | |
| Richness – constrained best 3 | Tillage – no till | -709.40 | 234.82 | | | | |
| Richness – constrained best 3 | Year | -265.52 | 215.28 | 1.52 | 1 | 0.23 | |

| **Model** | **Model term** | **Coefficient** | **Coefficient standard error** | **F** | **df** | ***P*-value** | **ΔAICc** |
|---|---|---|---|---|---|---|---|
| Richness – constrained best 3 | Field size (planted ha) | -0.26 | 2.70 | 0.01 | 1 | 0.93 | |
| Richness – constrained best 3 | Herbivore richness | -141.26 | 46.05 | 9.41 | 1 | 0.006 | |
| Richness – constrained best 3 | Parasitoid richness | 248.72 | 107.51 | 5.35 | 1 | 0.03 | |
| Richness – constrained best 3 | Pollinator richness | -29.94 | 21.14 | 2.01 | 1 | 0.17 | |
| Evenness – best | Intercept | 2451.24 | 181.95 | | 24 | | 0 |
| Evenness – best | Tillage – full till | 262.04 | 260.85 | 6.22 | 2 | 0.007 | |
| Evenness – best | Tillage – no till | -641.86 | 229.85 | | | | |
| Evenness – best | Year | -361.30 | 200.35 | 3.25 | 1 | 0.08 | |
| Evenness – constrained best | Intercept | 2434.26 | 211.27 | | 23 | | 0 |
| Evenness – constrained best | Tillage – full till | 280.47 | 287.83 | 4.74 | 2 | 0.02 | |
| Evenness – constrained best | Tillage – no till | -654.35 | 246.05 | | | | |
| Evenness – constrained best | Year | -370.64 | 211.90 | 3.06 | 1 | 0.09 | |
| Evenness – constrained best | Field size (planted ha) | 0.51 | 3.04 | 0.03 | 1 | 0.87 | |

**Table S9:** Test statistics for post-hoc Tukey tests of pairwise differences among tillage levels, for models where tillage was important. When the impact of tillage on biodiversity was assessed via negative binomial regression (abundance) we report Z values. When it was assessed via general linear models with the identity link (richness), we report t values. Shading separates models.

| **Functional group** | **Metric** | **Model #** | **Comparison** | **Z/t** | ***P*-value** |
|---|---|---|---|---|---|
| Kleptoparasite | Abundance | 3 | Intermediate vs. full | 2.50 | 0.03 |
| Kleptoparasite | Abundance | 3 | Intermediate vs. no | -0.50 | 0.87 |
| Kleptoparasite | Abundance | 3 | Full vs. no | -2.60 | 0.03 |
| Kleptoparasite | Abundance | 4, constrained 1 | Intermediate vs. full | 2.14 | 0.08 |
| Kleptoparasite | Abundance | 4, constrained 1 | Intermediate vs. no | -0.27 | 0.96 |
| Kleptoparasite | Abundance | 4, constrained 1 | Full vs. no | -2.10 | 0.09 |
| Pollinator | Richness | 1 | Intermediate vs. full | 0.80 | 0.71 |
| Pollinator | Richness | 1 | Intermediate vs. no | 3.34 | 0.01 |
| Pollinator | Richness | 1 | Full vs. no | 1.73 | 0.22 |
| Pollinator | Richness | 2 | Intermediate vs. full | 1.42 | 0.35 |
| Pollinator | Richness | 2 | Intermediate vs. no | 2.86 | 0.02 |
| Pollinator | Richness | 2 | Full vs. no | 0.93 | 0.63 |
| Pollinator | Richness | 3, constrained 1 | Intermediate vs. full | 0.82 | 0.69 |
| Pollinator | Richness | 3, constrained 1 | Intermediate vs. no | 3.20 | 0.01 |
| Pollinator | Richness | 3, constrained 1 | Full vs. no | 1.55 | 0.28 |
| Pollinator | Richness | Constrained 2 | Intermediate vs. full | 0.46 | 0.89 |
| Pollinator | Richness | Constrained 2 | Intermediate vs. no | 3.49 | 0.005 |
| Pollinator | Richness | Constrained 2 | Full vs. no | 2.00 | 0.13 |
| Crop yield | Abundance | 1 | Intermediate vs. full | -1.01 | 0.58 |
| Crop yield | Abundance | 1 | Intermediate vs. no | 2.83 | 0.02 |
| Crop yield | Abundance | 1 | Full vs. no | 3.27 | 0.01 |
| Crop yield | Abundance | 2 | Intermediate vs. full | -1.22 | 0.45 |
| Crop yield | Abundance | 2 | Intermediate vs. no | 2.76 | 0.03 |
| Crop yield | Abundance | 2 | Full vs. no | 3.42 | 0.01 |
| Crop yield | Abundance | 3 | Intermediate vs. full | -1.10 | 0.52 |
| Crop yield | Abundance | 3 | Intermediate vs. no | 2.58 | 0.04 |
| Crop yield | Abundance | 3 | Full vs. no | 3.19 | 0.01 |

| **Functional group** | **Metric** | **Model #** | **Comparison** | **Z/t** | ***P*-value** |
|---|---|---|---|---|---|
| Crop yield | Abundance | Constrained 1 | Intermediate vs. full | -1.06 | 0.55 |
| Crop yield | Abundance | Constrained 1 | Intermediate vs. no | 2.75 | 0.03 |
| Crop yield | Abundance | Constrained 1 | Full vs. no | 2.87 | 0.02 |
| Crop yield | Abundance | Constrained 2 | Intermediate vs. full | -1.31 | 0.41 |
| Crop yield | Abundance | Constrained 2 | Intermediate vs. no | 2.74 | 0.03 |
| Crop yield | Abundance | Constrained 2 | Full vs. no | 3.07 | 0.01 |
| Crop yield | Richness | 1 | Intermediate vs. full | -1.73 | 0.21 |
| Crop yield | Richness | 1 | Intermediate vs. no | 2.97 | 0.02 |
| Crop yield | Richness | 1 | Full vs. no | 4.12 | 0.001 |
| Crop yield | Richness | 2 | Intermediate vs. full | -1.43 | 0.34 |
| Crop yield | Richness | 2 | Intermediate vs. no | 3.39 | 0.01 |
| Crop yield | Richness | 2 | Full vs. no | 4.46 | <0.001 |
| Crop yield | Richness | 3 | Intermediate vs. full | -1.24 | 0.44 |
| Crop yield | Richness | 3 | Intermediate vs. no | 3.07 | 0.01 |
| Crop yield | Richness | 3 | Full vs. no | 3.68 | 0.003 |
| Crop yield | Richness | Constrained 1 | Intermediate vs. full | -1.45 | 0.33 |
| Crop yield | Richness | Constrained 1 | Intermediate vs. no | 2.62 | 0.04 |
| Crop yield | Richness | Constrained 1 | Full vs. no | 3.15 | 0.01 |
| Crop yield | Richness | Constrained 2 | Intermediate vs. full | -1.03 | 0.57 |
| Crop yield | Richness | Constrained 2 | Intermediate vs. no | 2.74 | 0.03 |
| Crop yield | Richness | Constrained 2 | Full vs. no | 2.83 | 0.02 |
| Crop yield | Richness | Constrained 3 | Intermediate vs. full | -1.30 | 0.41 |
| Crop yield | Richness | Constrained 3 | Intermediate vs. no | 3.02 | 0.02 |
| Crop yield | Richness | Constrained 3 | Full vs. no | 3.50 | 0.01 |
| Crop yield | Evenness | 1 | Intermediate vs. full | -1.00 | 0.58 |
| Crop yield | Evenness | 1 | Intermediate vs. no | 2.79 | 0.03 |
| Crop yield | Evenness | 1 | Full vs. no | 3.24 | 0.01 |
| Crop yield | Evenness | Constrained 1 | Intermediate vs. full | -0.97 | 0.60 |
| Crop yield | Evenness | Constrained 1 | Intermediate vs. no | 2.66 | 0.04 |
| Crop yield | Evenness | Constrained 1 | Full vs. no | 2.76 | 0.03 |

**Table S10:** Test statistics for multivariate analyses of local and landscape habitat on insect community composition. Non-categorical variables were converted to categories that separated sites below ("low") and above ("high") the median. Shading separates models.

| **Functional group** | **Model term** | **F** | **df** | ***P*-value** |
|---|---|---|---|---|
| Herbivore | Tillage | 0.13 | 2 | 0.98 |
| Herbivore | Proportion semi-natural habitat | 1.50 | 1 | 0.24 |
| Herbivore | Tillage – semi-natural habitat interaction | 0.01 | 2 | 0.99 |
| Herbivore | Year | 0.68 | 1 | 0.60 |
| Herbivore | Degree days before sampling | 0.38 | 1 | 0.79 |
| Herbivore | Cumulative precipitation before sampling | 2.06 | 1 | 0.11 |
| Herbivore | Field size (ha planted) | 0.94 | 1 | 0.47 |
| Herbivore | Residual | | 20 | |
| Pollinator | Tillage | 1.02 | 2 | 0.46 |
| Pollinator | Proportion semi-natural habitat | 1.48 | 1 | 0.14 |
| Pollinator | Tillage – semi-natural habitat interaction | 1.04 | 2 | 0.42 |
| Pollinator | Year | 1.79 | 1 | 0.06 |
| Pollinator | Degree days before sampling | 0.60 | 1 | 0.84 |
| Pollinator | Cumulative precipitation before sampling | 0.55 | 1 | 0.87 |
| Pollinator | Field size (ha planted) | 0.14 | 1 | 0.99 |
| Pollinator | Residual | | 20 | |
| Predator | Tillage | 1.72 | 2 | 0.053 |
| Predator | Proportion semi-natural habitat | 0.93 | 1 | 0.51 |

| **Functional group** | **Model term** | **F** | **df** | ***P*-value** |
|---|---|---|---|---|
| Predator | Tillage – semi-natural habitat interaction | 0.72 | 2 | 0.76 |
| Predator | Year | 1.98 | 1 | 0.06 |
| Predator | Degree days before sampling | 1.72 | 1 | 0.11 |
| Predator | Cumulative precipitation before sampling | 0.39 | 1 | 0.89 |
| Predator | Field size (ha planted) | 1.13 | 1 | 0.37 |
| Predator | Residual | | 20 | |

**Table S11:** Correlations among abundance and diversity measures, within and across functional groups.

| **Metric 1** | **Metric 2** | **Test statistic (S)** | **r** | ***P*-value** |
|---|---|---|---|---|
| Natural enemy abundance | Natural enemy evenness | 8502.89 | -0.89 | <0.0001 |
| Natural enemy abundance | Herbivore evenness | 7318.63 | -0.63 | 0.0002 |
| Natural enemy abundance | Pollinator evenness | 6508.45 | -0.45 | 0.0131 |
| Natural enemy abundance | Predator evenness | 5643.54 | -0.54 | 0.0027 |
| Natural enemy abundance | Herbivore abundance | 964.21 | 0.79 | <0.0001 |
| Natural enemy abundance | Pollinator abundance | 2218.49 | 0.51 | 0.0043 |
| Natural enemy abundance | Predator abundance | 1270.91 | 0.72 | <0.0001 |
| Natural enemy abundance | Natural enemy richness | 1142.54 | 0.75 | <0.0001 |
| Natural enemy abundance | Herbivore richness | 2370.79 | 0.47 | 0.0084 |
| Natural enemy abundance | Kleptoparasite richness | 2618.28 | 0.42 | 0.0217 |
| Natural enemy abundance | Parasitoid richness | 2083.54 | 0.54 | 0.0022 |
| Natural enemy abundance | Pollinator richness | 2824.79 | 0.37 | 0.0432 |
| Natural enemy abundance | Predator richness | 1492.93 | 0.67 | 0.0001 |
| Herbivore abundance | Natural enemy evenness | 8114.00 | -0.81 | <0.0001 |
| Herbivore abundance | Herbivore evenness | 8074.00 | -0.80 | <0.0001 |
| Herbivore abundance | Pollinator evenness | 6022.00 | -0.34 | 0.0668 |
| Herbivore abundance | Predator evenness | 5073.19 | -0.39 | 0.0411 |
| Herbivore abundance | Kleptoparasite abundance | 3103.84 | 0.31 | 0.0961 |
| Herbivore abundance | Parasitoid abundance | 705.05 | 0.84 | <0.0001 |
| Herbivore abundance | Pollinator abundance | 2428.00 | 0.46 | 0.0113 |
| Herbivore abundance | Predator abundance | 2291.04 | 0.49 | 0.0059 |
| Herbivore abundance | Natural enemy richness | 2131.28 | 0.53 | 0.0028 |
| Herbivore abundance | Herbivore richness | 1940.23 | 0.57 | 0.0011 |
| Herbivore abundance | Kleptoparasite richness | 3160.45 | 0.30 | 0.1111 |
| Herbivore abundance | Parasitoid richness | 2360.77 | 0.47 | 0.0080 |

| Metric 1 | Metric 2 | Test statistic (S) | r | *P*-value |
|---|---|---:|---:|---:|
| Herbivore abundance | Pollinator richness | 3114.00 | 0.31 | 0.0986 |
| Herbivore abundance | Predator richness | 2465.99 | 0.45 | 0.0123 |
| Kleptoparasite abundance | Natural enemy evenness | 5862.00 | -0.30 | 0.1023 |
| Kleptoparasite abundance | Herbivore evenness | 5240.91 | -0.17 | 0.3808 |
| Kleptoparasite abundance | Pollinator evenness | 7094.11 | -0.58 | 0.0008 |
| Kleptoparasite abundance | Predator evenness | 4363.59 | -0.19 | 0.3221 |
| Kleptoparasite abundance | Parasitoid abundance | 2867.52 | 0.36 | 0.0493 |
| Kleptoparasite abundance | Pollinator abundance | 1559.67 | 0.65 | 0.0001 |
| Kleptoparasite abundance | Predator abundance | 2768.61 | 0.38 | 0.0361 |
| Kleptoparasite abundance | Natural enemy richness | 1794.43 | 0.60 | 0.0004 |
| Kleptoparasite abundance | Herbivore richness | 4514.40 | 0.00 | 0.9819 |
| Kleptoparasite abundance | Kleptoparasite richness | 150.19 | 0.97 | <0.0001 |
| Kleptoparasite abundance | Parasitoid richness | 3044.58 | 0.32 | 0.0820 |
| Kleptoparasite abundance | Pollinator richness | 2733.33 | 0.39 | 0.0322 |
| Kleptoparasite abundance | Predator richness | 2035.67 | 0.55 | 0.0018 |
| Parasitoid abundance | Natural enemy evenness | 8493.11 | -0.89 | <0.0001 |
| Parasitoid abundance | Herbivore evenness | 7883.64 | -0.75 | <0.0001 |
| Parasitoid abundance | Pollinator evenness | 6333.43 | -0.41 | 0.0248 |
| Parasitoid abundance | Predator evenness | 4558.87 | -0.25 | 0.2039 |
| Parasitoid abundance | Pollinator abundance | 2450.41 | 0.45 | 0.0116 |
| Parasitoid abundance | Predator abundance | 2548.75 | 0.43 | 0.0169 |
| Parasitoid abundance | Natural enemy richness | 1674.71 | 0.63 | 0.0002 |
| Parasitoid abundance | Herbivore richness | 2739.06 | 0.39 | 0.0328 |
| Parasitoid abundance | Kleptoparasite richness | 3034.29 | 0.32 | 0.0797 |
| Parasitoid abundance | Parasitoid richness | 2010.28 | 0.55 | 0.0015 |
| Parasitoid abundance | Pollinator richness | 2913.69 | 0.35 | 0.0566 |
| Parasitoid abundance | Predator richness | 2091.10 | 0.53 | 0.0023 |
| Pollinator abundance | Natural enemy evenness | 6670.00 | -0.48 | 0.0073 |
| Pollinator abundance | Herbivore evenness | 5880.00 | -0.31 | 0.0979 |
| Pollinator abundance | Pollinator evenness | 8582.00 | -0.91 | <0.0001 |
| Pollinator abundance | Predator evenness | 5060.19 | -0.38 | 0.0432 |
| Pollinator abundance | Predator abundance | 2430.16 | 0.46 | 0.0107 |
| Pollinator abundance | Natural enemy richness | 2216.09 | 0.51 | 0.0042 |
| Pollinator abundance | Herbivore richness | 4294.19 | 0.04 | 0.8147 |
| Pollinator abundance | Kleptoparasite richness | 1292.28 | 0.71 | <0.0001 |
| Pollinator abundance | Parasitoid richness | 2790.84 | 0.38 | 0.0388 |
| Pollinator abundance | Pollinator richness | 1029.96 | 0.77 | <0.0001 |

| **Metric 1** | **Metric 2** | **Test statistic (S)** | **r** | ***P*-value** |
|---|---|---:|---:|---:|
| Pollinator abundance | Predator richness | 2497.45 | 0.44 | 0.0139 |
| Predator abundance | Natural enemy evenness | 6819.07 | -0.52 | 0.0034 |
| Predator abundance | Herbivore evenness | 6158.48 | -0.37 | 0.0441 |
| Predator abundance | Pollinator evenness | 6352.65 | -0.41 | 0.0232 |
| Predator abundance | Predator evenness | 6827.41 | -0.87 | <0.0001 |
| Predator abundance | Natural enemy richness | 1508.16 | 0.66 | 0.0001 |
| Predator abundance | Herbivore richness | 1956.22 | 0.56 | 0.0011 |
| Predator abundance | Kleptoparasite richness | 2922.22 | 0.35 | 0.0580 |
| Predator abundance | Parasitoid richness | 3210.10 | 0.29 | 0.1257 |
| Predator abundance | Pollinator richness | 3065.09 | 0.32 | 0.0867 |
| Predator abundance | Predator richness | 1510.23 | 0.66 | 0.0001 |
| Natural enemy richness | Herbivore richness | 2368.06 | 0.47 | 0.0083 |
| Natural enemy richness | Kleptoparasite richness | 1877.48 | 0.58 | 0.0007 |
| Natural enemy richness | Parasitoid richness | 1976.75 | 0.56 | 0.0013 |
| Natural enemy richness | Pollinator richness | 2838.83 | 0.37 | 0.0451 |
| Natural enemy richness | Predator richness | 304.32 | 0.93 | <0.0001 |
| Herbivore richness | Kleptoparasite richness | 4487.27 | 0.00 | 0.9928 |
| Herbivore richness | Parasitoid richness | 3507.91 | 0.22 | 0.2436 |
| Herbivore richness | Pollinator richness | 4586.03 | -0.02 | 0.9154 |
| Herbivore richness | Predator richness | 2195.77 | 0.51 | 0.0039 |
| Kleptoparasite richness | Parasitoid richness | 3032.66 | 0.33 | 0.0794 |
| Kleptoparasite richness | Pollinator richness | 2545.46 | 0.43 | 0.0166 |
| Kleptoparasite richness | Predator richness | 2057.67 | 0.54 | 0.0020 |
| Parasitoid richness | Pollinator richness | 2990.25 | 0.33 | 0.0706 |
| Parasitoid richness | Predator richness | 3355.26 | 0.25 | 0.1764 |
| Pollinator richness | Predator richness | 3203.21 | 0.29 | 0.1236 |
| Natural enemy evenness | Herbivore evenness | 1596.00 | 0.64 | 0.0002 |
| Natural enemy evenness | Pollinator evenness | 2586.00 | 0.42 | 0.0201 |
| Natural enemy evenness | Predator evenness | 1667.73 | 0.54 | 0.0028 |
| Natural enemy evenness | Natural enemy richness | 6516.43 | -0.45 | 0.0127 |
| Natural enemy evenness | Herbivore richness | 6215.09 | -0.38 | 0.0369 |
| Natural enemy evenness | Kleptoparasite richness | 5780.76 | -0.29 | 0.1254 |
| Natural enemy evenness | Parasitoid richness | 6264.59 | -0.39 | 0.0314 |
| Natural enemy evenness | Pollinator richness | 6116.70 | -0.36 | 0.0502 |
| Natural enemy evenness | Predator richness | 6362.63 | -0.42 | 0.0224 |
| Herbivore evenness | Pollinator evenness | 3462.00 | 0.23 | 0.2209 |
| Herbivore evenness | Predator evenness | 3067.92 | 0.16 | 0.4149 |

| **Metric 1** | **Metric 2** | **Test statistic (S)** | **r** | ***P*-value** |
|---|---|---|---|---|
| Herbivore evenness | Natural enemy richness | 6476.04 | -0.44 | 0.0148 |
| Herbivore evenness | Herbivore richness | 6154.23 | -0.37 | 0.0447 |
| Herbivore evenness | Kleptoparasite richness | 5206.49 | -0.16 | 0.4035 |
| Herbivore evenness | Parasitoid richness | 5356.20 | -0.19 | 0.3105 |
| Herbivore evenness | Pollinator richness | 5651.35 | -0.26 | 0.1699 |
| Herbivore evenness | Predator richness | 6559.54 | -0.46 | 0.0107 |
| Pollinator evenness | Predator evenness | 2395.83 | 0.34 | 0.0728 |
| Pollinator evenness | Natural enemy richness | 6570.95 | -0.46 | 0.0102 |
| Pollinator evenness | Herbivore richness | 4444.29 | 0.01 | 0.9528 |
| Pollinator evenness | Kleptoparasite richness | 7374.50 | -0.64 | 0.0001 |
| Pollinator evenness | Parasitoid richness | 5873.13 | -0.31 | 0.0994 |
| Pollinator evenness | Pollinator richness | 6886.93 | -0.53 | 0.0025 |
| Pollinator evenness | Predator richness | 6467.17 | -0.44 | 0.0153 |
| Predator evenness | Natural enemy richness | 4482.30 | -0.23 | 0.2461 |
| Predator evenness | Herbivore richness | 4969.25 | -0.36 | 0.0599 |
| Predator evenness | Kleptoparasite richness | 4238.79 | -0.16 | 0.4159 |
| Predator evenness | Parasitoid richness | 4185.16 | -0.15 | 0.4605 |
| Predator evenness | Pollinator richness | 4461.04 | -0.22 | 0.2587 |
| Predator evenness | Predator richness | 4527.54 | -0.24 | 0.2205 |

**Table S12** is in a separate Excel file. It shows abundance of each herbivore at each site.

**Figure S1:** Map of study sites in 2013 (circles) and 2014 (triangles). Colors indicate whether sites were subject to full (red), intermediate (yellow), or no (blue) tillage. The vertical line between Palouse and Moscow is the Washington-Idaho state line.

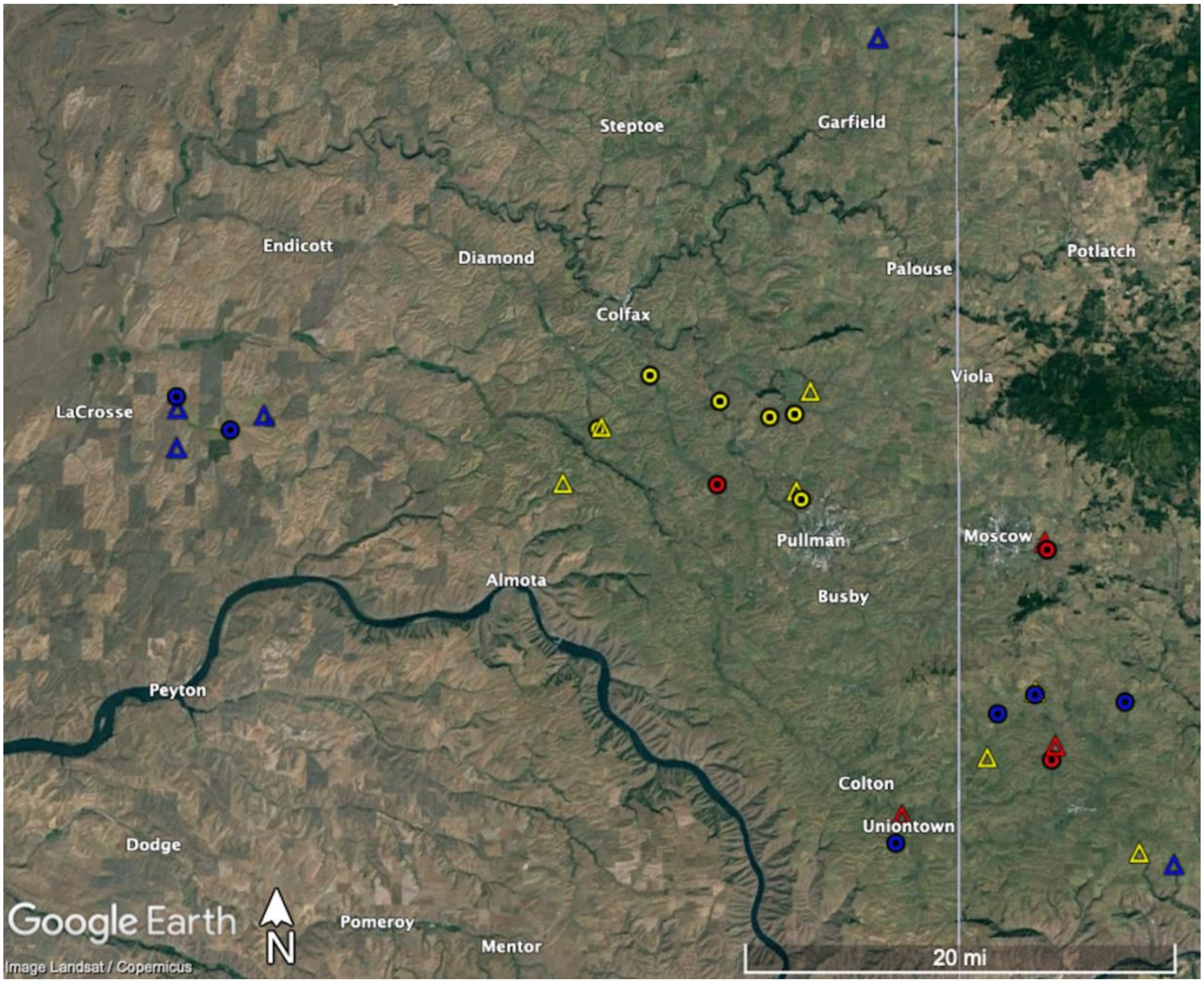

**Figure S2:** Box and whisker plot of field size by tillage regime. “Int.” is intermediate tillage. Boxes span the $25^{th}$ to $75^{th}$ percentiles, whiskers extend 1.5 times the interquartile range beyond the boxes, and there is a line at the median. Lines with asterisks indicate groups that are statistically different from each other at $\alpha = 0.05$.

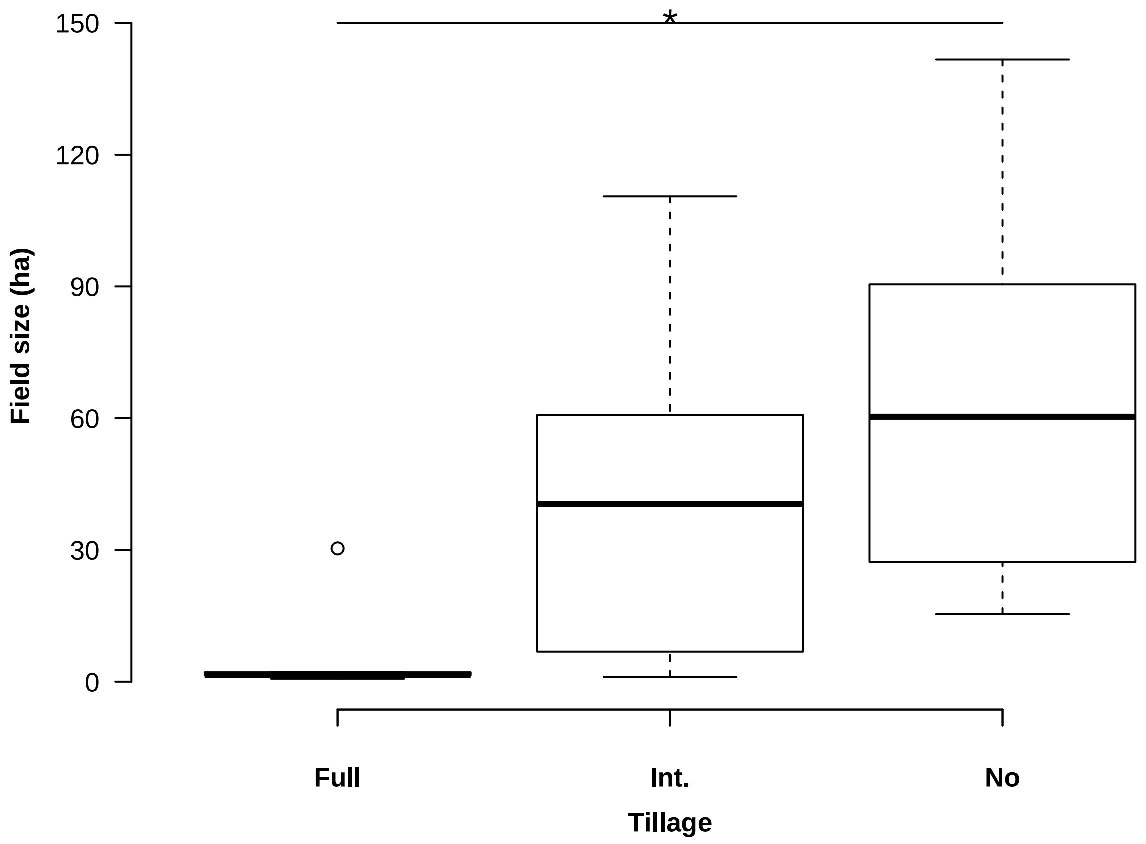

**Figure S3:** Principle component analysis plots showing how herbivore (a, b), pollinator (c, d), and predator (e, f) communities vary with tillage (a, c, e) and the proportion of semi-natural habitat in the landscape (b, d, f). Red indicates full tillage, black intermediate tillage, and blue no tillage. Green indicates low amounts of semi-natural habitat and orange indicates high. Ellipses show standard deviations of the data.

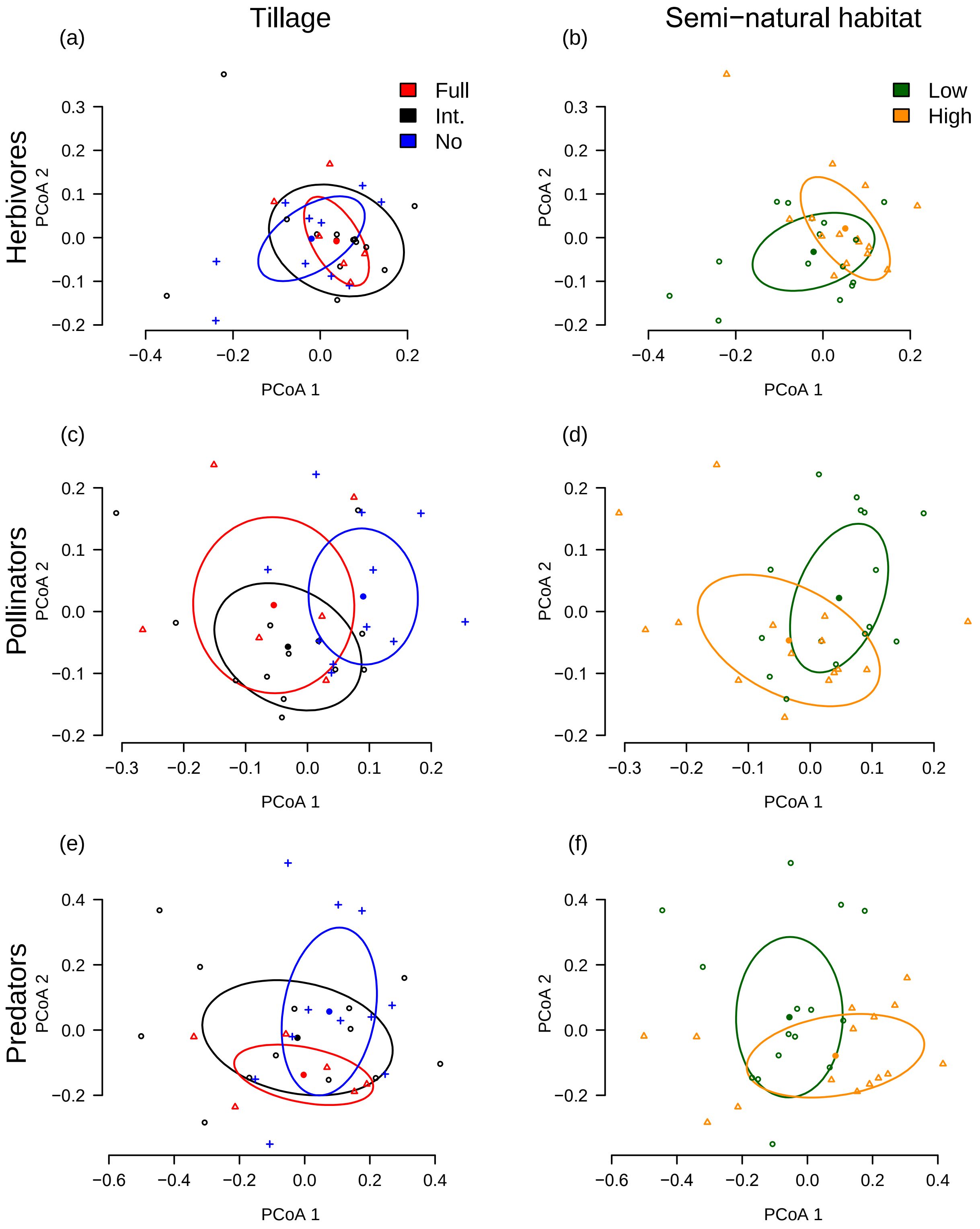
